# PNPO-PLP Axis Senses Prolonged Hypoxia by Regulating Lysosomal Activity

**DOI:** 10.1101/2022.10.28.514185

**Authors:** Hiroki Sekine, Haruna Takeda, Akihiro Kishino, Hayato Anzawa, Takayuki Isagawa, Nao Ohta, Nobufumi Kato, Shu Kimura, Zun Liu, Koichiro Kato, Fumiki Katsuoka, Masayuki Yamamoto, Fumihito Miura, Takashi Ito, Masatomo Takahashi, Yoshihiro Izumi, Hiroyuki Fujita, Hitoshi Yamagata, Norihiko Takeda, Takeshi Bamba, Norio Suzuki, Kengo Kinoshita, Hozumi Motohashi

## Abstract

Oxygen is critical for all metazoan organisms on the earth and impacts various biological processes in physiological and pathological conditions. While oxygen sensing systems inducing acute hypoxic response, including HIF pathway, have been identified, those operating in prolonged hypoxia remain to be elucidated. Here, we show that pyridoxine 5’-phosphate oxidase (PNPO) that catalyzes bioactivation of vitamin B6 serves as an oxygen sensor and regulates lysosomal activity in macrophages. Decline of PNPO activity under prolonged hypoxia reduced an active form of vitamin B6, pyridoxal 5’-phosphate (PLP), and inhibited lysosomal activity, leading to the augmentation of inflammatory response of macrophages. The PNPO-PLP axis creates a distinct layer of oxygen sensing, which gradually turns down and up the PLP-dependent metabolism according to prolonged changes in oxygen availability.

## Introduction

Oxygenation of the Earth’s atmosphere resulted in the emergence of aerobic organisms that utilize molecular oxygen (O_2_) for energy metabolism in mitochondria and many other reactions catalyzed by oxygenases and oxidases. Oxygenases incorporate oxygen atoms from O_2_ into their substrates whereas oxidases use O_2_ as an electron acceptor. All metazoan organisms have been shown to possess oxygen-sensing mechanisms for adaptation to low oxygen conditions^1^, which utilize dioxygenases with high K_m_ values for oxygen, namely PHD^2^, KDM6A^3^ and ADO^4^, as oxygen sensors. However, how oxidases make contributions to oxygen sensing and response to hypoxia is not fully understood.

Among the dioxygenases, PHD and its effector HIF form a major molecular system that mediates the response to hypoxia^5^. PHD belongs to the 2-oxoglutarate (2OG)-dependent dioxygenase family and requires molecular oxygen as a substrate^6,7^. Because the K_m_ value of PHD for oxygen is high enough, PHD activity easily declines under hypoxia, resulting in the stabilization of HIFα subunits and activation of HIF target genes. Thus, a high K_m_ value for oxygen allows PHD to serve as an oxygen sensor. KDM6A has been shown to serve as another oxygen sensor, regulating the epigenetic status in response to hypoxia, because the K_m_ value of KDM6A for oxygen is similarly high to that of PHD^2,3^. In contrast, KDM5A and TET activities have been reported to be sensitive to oxygen tension, although their K_m_ values for O_2_ are rather low^3,8-10^, implying alternative oxygen-sensing mechanisms for the regulation of 2OG-dependent dioxygenase activities.

Crosstalk between hypoxia and inflammation is understood at the molecular level as a functional interaction between HIF-1α, a key regulator of the hypoxic response, and NF-κB, a key regulator of the inflammatory response^11^. HIF-1α has been shown to enhance IL-1β production and drive inflammation by inducing a metabolic shift to glycolysis from oxidative phosphorylation in activated macrophages^12^. Deleting HIF-1α in myeloid cells attenuates inflammation, verifying an important role of HIF-1α as a proinflammatory regulator *in vivo*^13^. In good agreement with the HIF-1α requirement in the inflammatory response of macrophages, acute hypoxia augments the inflammatory response by activating NF-κB signaling *in vitro* and in mice^11^. These studies provide solid evidence for the proinflammatory effects of acute hypoxia, which is mediated by HIF-1α. However, this mechanistic scheme appears less applicable to prolonged hypoxia because of the transient nature of HIF-1α-mediated transcriptional activation^11^. A question here is how the prolonged hypoxia is sensed, responded, and involved in the inflammatory processes.

Pyridoxal 5’-phosphate (PLP) is an active form of vitamin B6 and serves as a coenzyme for many amino acid-metabolizing enzymes. Pyridoxine that is a major form of vitamin B6 in food undergoes bioactivation to become PLP, which is catalyzed by pyridoxine 5’-phosphate oxidase (PNPO) requiring molecular oxygen as a substrate. Here, we demonstrate that PNPO serves as an oxygen sensor and regulates lysosomal activity under prolonged hypoxia, resulting in exacerbated inflammation, irrespective of the HIF pathway status. Intriguingly, prolonged hypoxia, but not acute hypoxia, reduces lysosomal activity in macrophages, which limits ferrous iron (Fe^2+^) availability and switches off TET2 function that mediates inflammatory resolution. Metabolome analysis revealed that prolonged hypoxia suppresses vitamin B6 bioactivation catalyzed by PNPO. Supplementation of active vitamin B6 restored lysosomal function and TET2 protein accumulation in macrophages and attenuated inflammatory response in mice under prolonged hypoxia. This study has identified PNPO-PLP axis as an alternative oxygen-sensing system operative in prolonged hypoxia, which is distinct from dioxygenase-dependent mechanisms including PHD-HIF pathway.

## Results

### Prolonged hypoxia exacerbates inflammation

To examine the impact of prolonged hypoxia on inflammation, we used a genetically engineered mouse model of systemic hypoxia, inherited super-anemic mice (ISAM), which exhibit severe anemia caused by erythropoietin insufficiency and consequent tissue hypoxia^14,15^. ISAM and control mice were subjected to dextran sulfate sodium (DSS)-induced colitis. ISAM were more susceptible to DSS-induced colitis, showing significant exacerbation of body weight loss (Fig. 1a), colon shortening (Fig. 1b and 1c) and histopathological damage (Fig. 1d and 1e). The expression of the proinflammatory cytokine genes, *Il6, Il23*, and *Il1b*, in colon tissues was higher in ISAM than in control mice with DSS treatment, although *Tnfa* and *Il12b* were similarly increased in the colon tissues of both mouse lines (Fig. 1f). To explore the possibility that these phenotypes in ISAM were attributed to the enhanced proinflammatory response of macrophages, which play a key role in DSS-induced colitis, we collected peritoneal macrophages and examined their response to LPS stimulation. The proinflammatory cytokine genes *Il6* and *Il1b* and the anti-inflammatory cytokine gene *Il10* were higher and lower, respectively, in peritoneal macrophages from ISAM than in those from control mice (Fig. 1g). Consistently, peritoneal macrophages of ISAM secreted increased amounts of IL-6 and IL-1β (Fig. 1h). These results suggested that ISAM macrophages were more proinflammatory than control macrophages. To examine whether the proinflammatory phenotypes of ISAM macrophages were acquired due to a hypoxic environment or their intrinsic properties, we cultured bone marrow cells from ISAM and control mice under normoxia and obtained bone marrow-derived macrophages (BMDMs). The expression levels of *Il6, Il1b*, and *Il10* in response to LPS were all comparable between BMDMs derived from ISAM and control mice (Extended Data Fig. 1), suggesting that a prolonged hypoxic environment promoted the proinflammatory phenotypes of ISAM macrophages.

**Fig. 1:**
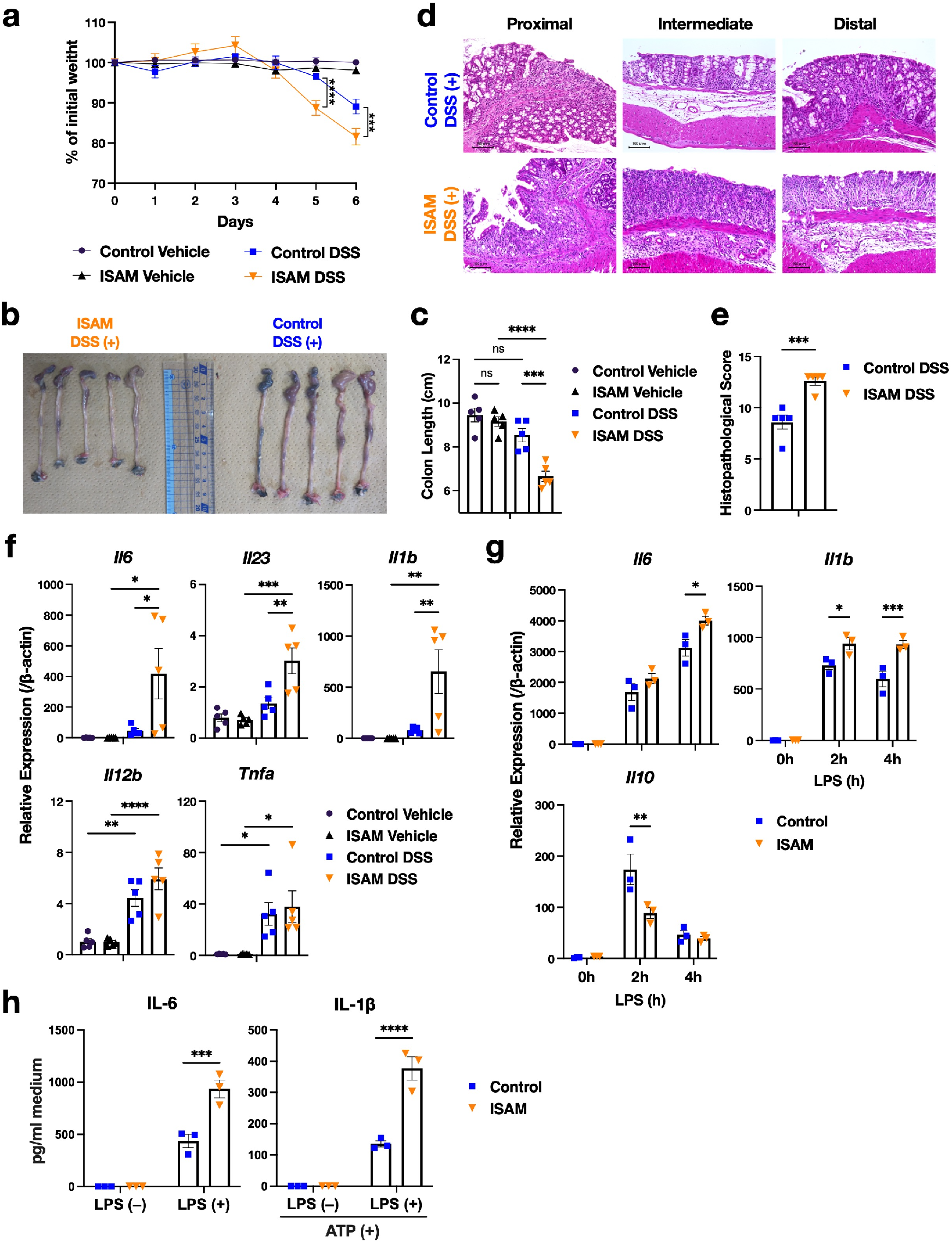
Prolonged hypoxia model mice ISAM are highly susceptible to DSS-induced colitis compared with control mice. **a**. Body weight changes in ISAM and control mice during treatment with 3% DSS or vehicle (water) (n = 5). **b-e**. Colonic pathological changes on day 6 of treatment with 3% DSS. Macroscopic appearance (b), length (c), HE-stained sections (d) and histopathological scores (e) of colons in ISAM and control mice (n = 5). **f**.Expression of proinflammatory cytokine genes in the colons of ISAM and control mice with or without DSS treatment for 6 days (n = 5). **g**. Expression of cytokine genes in peritoneal macrophages from ISAM and control mice after LPS treatment (n = 3). **h**. ELISA of cytokines in the culture supernatant of peritoneal macrophages from ISAM and control mice. Macrophages were stimulated with or without LPS for 12 h (n = 3). For IL-1β detection, additional incubation was conducted for 2 h in the presence of 1 mM ATP. Scale bars correspond to 100 μm (d). Error bars represent S.E.M. of 3-5 biological replicates. Two-way ANOVA (a, g, h), one-way ANOVA (c, f) and Student’s *t*-test (e) were conducted to evaluate statistical significance. *P < 0.05, **P < 0.01, ***P < 0.001, ****P < 0.0001.

### Macrophages differentiated under hypoxia acquire proinflammatory phenotypes

To verify the impact of oxygen tension on the inflammatory phenotypes of macrophages, we prepared BMDMs differentiated under normoxia and 1% oxygen, which were designated normoxia-BMDMs (Norm-BMDMs) and chronic hypoxia-BMDMs (CHyp-BMDMs), respectively (Extended Data Fig. 2a). Norm-BMDMs and CHyp-BMDMs were subsequently stimulated with lipopolysaccharide (LPS) under each respective oxygen tension. Acute hypoxia-BMDMs (AHyp-BMDMs) were differentiated under normoxia and stimulated with LPS under 1% oxygen (Extended Data Fig. 2a). Inflammatory responses of the three kinds of BMDMs were compared in terms of the LPS-induced transcriptome measured by RNA-seq analysis. Typical HIF target genes were upregulated both in AHyp-BMDMs and CHyp-BMDMs during LPS stimulation compared with Norm-BMDMs (Fig. 2a). Amazingly, remarkable upregulation and downregulation of proinflammatory and anti-inflammatory genes, respectively, were observed in CHyp-BMDMs but not in AHyp-BMDMs compared with Norm-BMDMs (Fig. 2a). Indeed, IL-6 and TNF-α were more abundantly secreted from CHyp-BMDMs than Norm-BMDMs in response to LPS (Extended Data Fig. 2b).

**Fig. 2:**
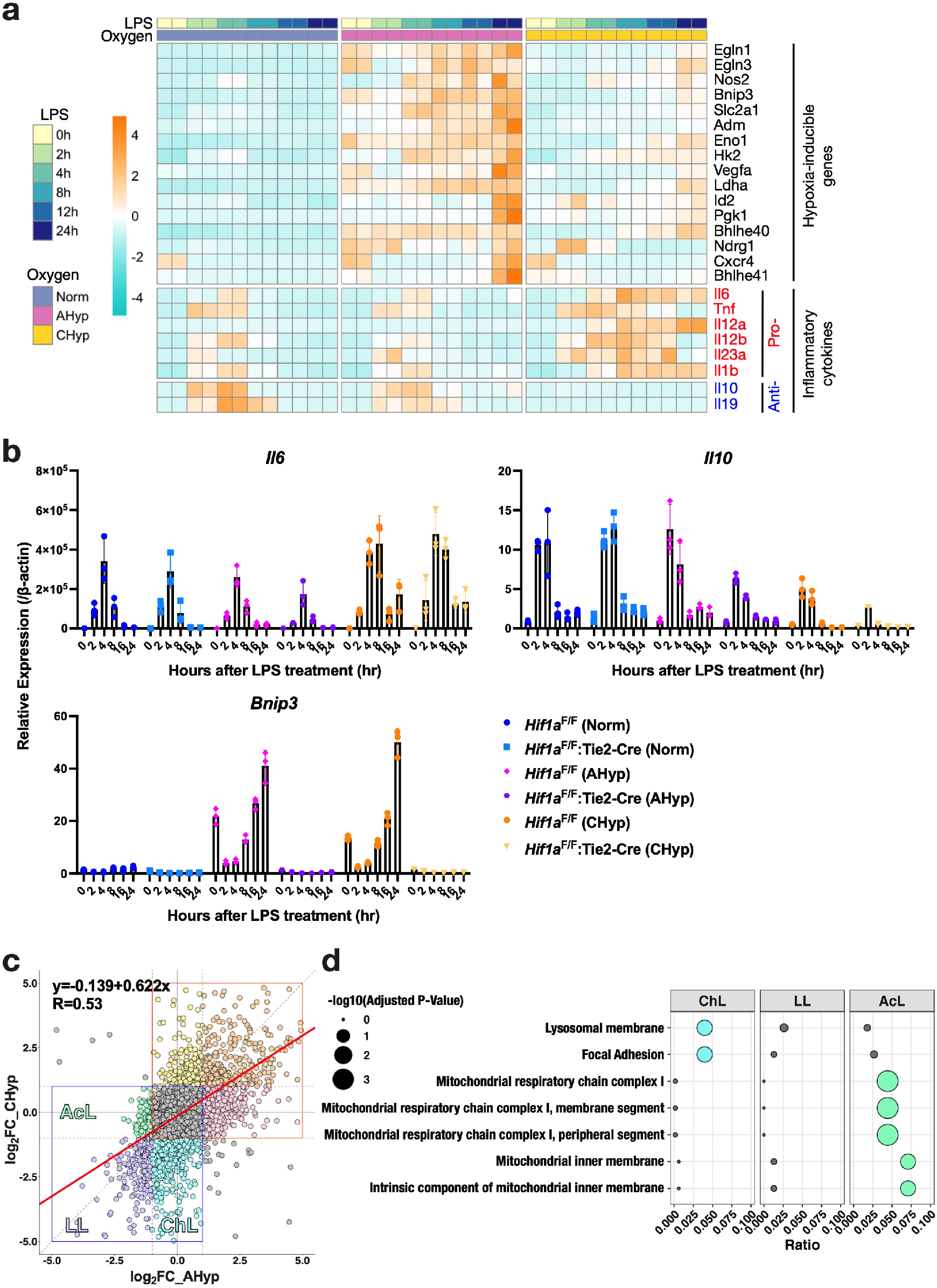
Prolonged hypoxia augments the proinflammatory response of BMDMs. **a**. A heatmap illustrating RNA-seq data of representative HIF target genes and cytokine genes in BMDMs after LPS stimulation. Norm: BMDMs differentiated and stimulated with LPS under normoxia, AHyp: BMDMs differentiated under normoxia and stimulated with LPS under 1% oxygen, CHyp: BMDMs differentiated and stimulated with LPS under 1% oxygen. **b**. Expression of cytokine genes, *Il6* and *Il10*, and a HIF target gene, *Bnip3*, in BMDMs after LPS stimulation (n = 3). Norm-, AHyp-, and CHyp-BMDMs generated from *Hif1a*^F/F^ mice and *Hif1a*^F/F^:Tie2-Cre mice were examined. Error bars represent S.E.M. of 3 biological replicates. **c**. Scatter plots showing a correlation of gene expression fold changes by prolonged and acute hypoxia in BMDMs. A horizontal axis indicates log_2_ fold change of CHyp vs. Norm, and a vertical axis indicates log_2_ fold change of AHyp vs. Norm. Areas enclosed by red and blue squares are those containing upregulated and downregulated genes, respectively. The strength of the correlation was evaluated with Pearson product-moment correlation coefficient. **d**. Enrichr analysis (GO_Cellular_Component_2017b) of downregulated genes. ChL, LL and AcL indicate gene groups specifically downregulated by prolonged hypoxia, commonly downregulated, and specifically downregulated by acute hypoxia, respectively (indicated in panel c). Circle sizes indicate adjusted P values, and circle colors indicate statistical significance (blue and green, adjusted P value < 0.05; gray, not significant).

We next examined whether HIF pathway is involved in the gene expression alteration in CHyp-BMDMs. A previous study described that HIF-1α and HIF-2α promote proinflammatory and anti-inflammatory polarization of macrophages, respectively^16^, raising the possibility of a skewed balance between HIF-1α and HIF-2α in CHyp-BMDMs. As HIF-2α was hardly detected in our experimental setting regardless of oxygen tension (Extended Data Fig. 2c), we compared LPS-induced expression of cytokine genes and HIF-target genes, *Bnip3* and *Pgk1*, in BMDMs of control (*Hif1a*^F/F^) mice and *Hif1a* knockout (*Hif1a*^F/F^:Tie2-Cre) mice (Fig. 2b and Extended Data Fig. 2d). *Hif1a* deficiency, which was verified by abrogated expression of *Bnip3* and *Pgk1*, did not alter the proinflammatory gene expression in CHyp-BMDMs. Thus, we considered that the proinflammatory phenotypes of BMDMs under prolonged hypoxia were independent of the PHD-HIF pathway.

For comprehensive understanding of how prolonged and acute hypoxia influence the LPS-induced gene expression, we approximated total amount of transcripts over time as the area under curve (AUC) calculated from mRNA level at each time point from 0 to 24 hours after LPS addition. AUC ratios of CHyp-BMDMs and AHyp-BMDMs vs. Norm-BMDMs were plotted (Fig. 2c and Extended Data Fig. 3a). While overall comparison showed a positive correlation between the impacts of chronic and acute hypoxia on the LPS-induced gene expression (R = 0.53; Fig. 2c and middle panel in Extended Data Fig. 3a), genes were categorized into 7 classes, specifically upregulated in chronic hypoxia (ChH), specifically upregulated in acute hypoxia (AcH), commonly upregulated in both conditions (HH), specifically downregulated in chronic hypoxia (ChL), specifically downregulated in acute hypoxia (AcL), commonly downregulated in both conditions (LL) and not changed (NC) (Fig. 2c and Extended Data Fig. 3a). As expected from *in vivo* and *ex vivo* results of ISAM and control mice, pro- and anti-inflammatory cytokine genes were found in ChH and ChL classes, respectively (right and left panels in Extended Data Fig. 3a). Pathway analysis of upregulated genes revealed that genes bound by HIF, SMRT and RELA were enriched in AcH and HH classes, whereas RELA-bound genes were an only significant pathway enriched in ChH class (Extended Data Fig. 3b). Among downregulated genes, mitochondria-related pathways were enriched in AcL class (Fig. 2d), which is consistent with a previous report describing that mitochondrial function is dysregulated in acute hypoxia^17^. Intriguingly, lysosome-related pathway was enriched in ChL class (Fig. 2d and Extended Data Fig. 3c), implying that lysosomal activity was inhibited in BMDMs exposed to prolonged hypoxia.

### Prolonged hypoxia inhibits lysosomal activity

Because decline of lysosomal activity appeared to accompany the proinflammatory phenotypes of BMDMs under prolonged hypoxia, we examined lysosomal acidification in Norm-BMDMs and CHyp-BMDMs. Fluorescence intensity indicating the lysosomal acidification was decreased in CHyp-BMDMs compared with Norm-BMDMs irrespective of LPS treatment (Fig. 3a). Consistently, CHyp-BMDMs exhibited reduction in the protein level of Lamp1, which is a lysosomal membrane protein (Fig. 3b). These results suggest that prolonged hypoxia inhibits lysosomal activity. *Hif1a* deficiency allowed lysosomal inhibition due to the prolonged hypoxia, indicating that HIF activity is dispensable for the lysosomal response to the prolonged hypoxia (Fig. 3c).

**Fig. 3:**
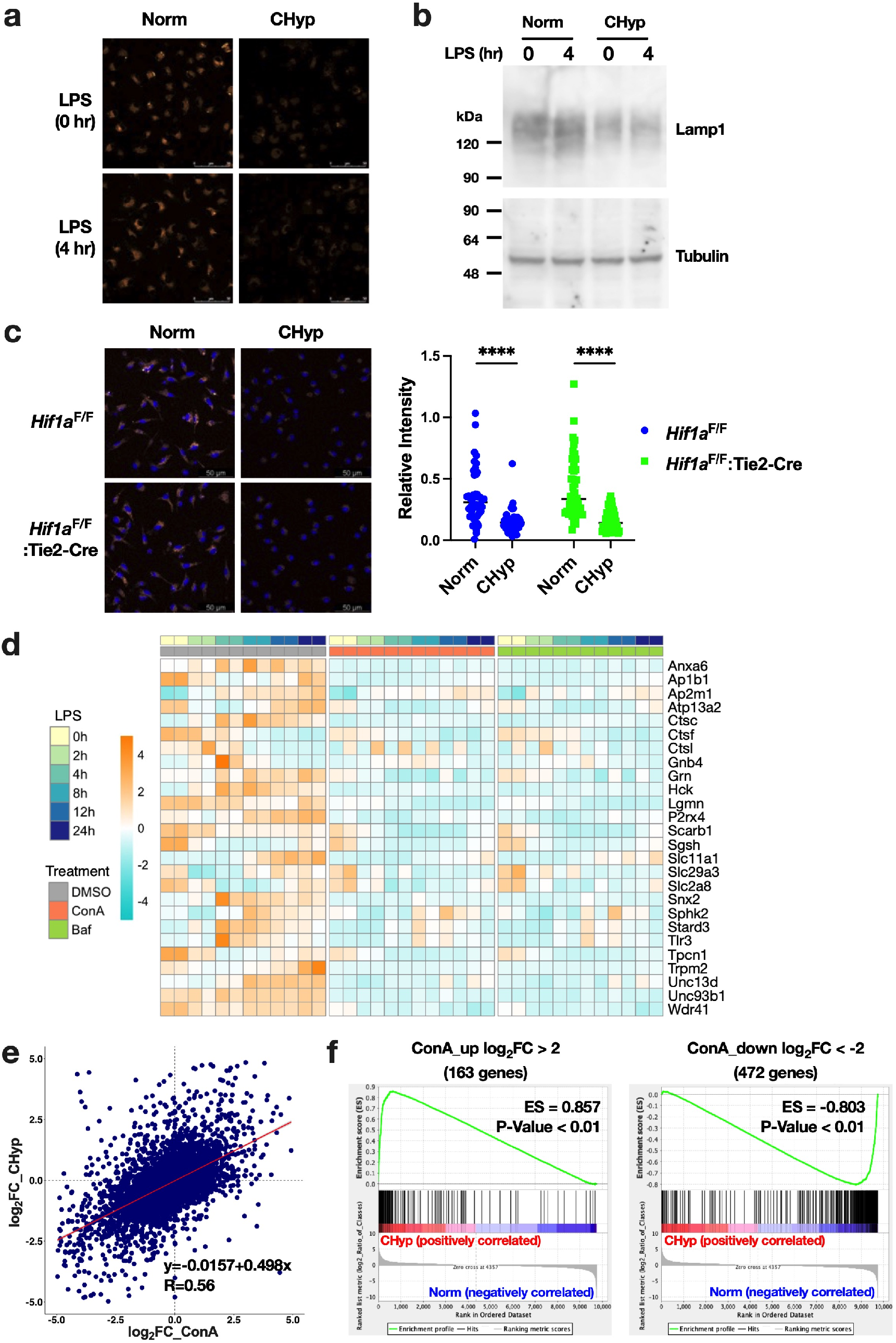
Lysosomal inhibition in prolonged hypoxia and impacts of lysosomal inhibition and prolonged hypoxia on the LPS-induced transcriptome. **a**. Representative AcidiFluor ORANGE staining for lysosomal acidification in BMDMs treated with or without LPS for 4 h. Norm, BMDMs differentiated and stimulated with LPS under normoxia; CHyp, BMDMs differentiated and stimulated with LPS under 1% oxygen. Scale bars correspond to 50 μm. **b**. Immunoblot analysis detecting Lamp1 in BMDMs treated with or without LPS for 4 h. Tubulin was detected as a loading control. **c**. Lysosomal acidification in BMDMs cultured from *Hif1a*^F/F^ and *Hif1a*^F/F^:Tie2-Cre mice. Norm: BMDMs differentiated under normoxia, CHyp: BMDMs differentiated under 1% oxygen. Scale bars correspond to 50 μm. Representative AcidiFluor ORANGE staining (left) and its quantification (right). Two-way ANOVA was conducted to evaluate statistical significance. ****P < 0.0001. **d**. A heatmap illustrating RNA-seq data of genes that belong to ChL class and assigned to “lysosome” in GO database, which is the same gene set shown in Extended Data Fig. 3c. BMDMs differentiated under normoxia were treated with 10 nM concanamycin A (ConA), 10 nM bafilomycin A (Baf) or vehicle (DMSO) at 16 h before LPS stimulation. **e**. Scatter plot showing a correlation of gene expression fold changes by prolonged hypoxia and lysosomal inhibition (ConA) in BMDMs. A horizontal axis indicates log_2_ fold change of ConA treatment vs. DMSO, and a vertical axis indicates log_2_ fold change of CHyp vs. Norm (shown in Fig. 2c). The strength of the correlation was evaluated with Pearson product-moment correlation coefficient. **f**. Gene set enrichment analysis comparing the impacts of prolonged hypoxia with lysosomal inhibition. Gene sets were defined as upregulated (left) or downregulated (right) genes by more than 4-fold (log_2_ 4) by ConA treatment. Changes in the LPS-induced transcriptome by prolonged hypoxia were analyzed against the gene sets.

We asked a question: to what extent lysosomal inhibition contributed to the transcriptome alteration under prolonged hypoxia. To compare impacts of lysosomal inhibition on the LPS-induced transcriptome with those of prolonged hypoxia, we conducted RNA-seq analysis of BMDMs differentiated under normoxia and treated with lysosomal inhibitors, concanamycin A (ConA) and bafilomycin A1 (Baf), or vehicle (DMSO) and examined how the lysosomal inhibitors influence the LPS-induced transcriptome. The lysosome-related pathway genes that were identified in the ChL class (see Extended Data Fig. 3c) were confirmed to decrease in both ConA-treated and Baf-treated BMDMs (Fig. 3d). We calculated AUC ratios of gene expression in ConA-treated and Baf-treated BMDMs vs. DMSO-treated BMDMs and verified that changes in the gene expression profiles induced by the two lysosomal inhibitors were highly matched (Extended Data Fig. 4a). The AUC ratios of ConA-treated and Baf-treated vs. DMSO-treated BMDMs were positively correlated with those of CHyp-BMDMs vs. Norm-BMDMs (Fig. 3e and Extended Data Fig. 4b). To further examine the similarity between them, Gene Set Enrichment Analysis (GSEA) was performed. Gene sets defined as upregulated and downregulated genes by ConA treatment were strongly enriched in the genes upregulated and downregulated by prolonged hypoxia, respectively (Fig. 3f). Similar results were obtained with gene sets defined by Baf treatment (Extended Data Fig. 4c and 4d). Conversely, gene sets defined as upregulated and downregulated genes by prolonged hypoxia were strongly enriched in the genes upregulated and downregulated by treatment with the lysosomal inhibitors, respectively (Extended Data Fig. 4e-4h). These results indicated a remarkable similarity between the effects of prolonged hypoxia and lysosomal inhibition on the LPS-induced transcriptional responses of macrophages, which let us suppose that lysosomal inhibition caused by prolonged hypoxia is responsible for proinflammatory phenotypes of macrophages.

### Prolonged hypoxia suppresses vitamin B6 bioactivation by PNPO

To explore the mechanism by which prolonged hypoxia suppresses lysosomal activity in BMDMs, we examined metabolome of Norm-BMDMs and CHyp-BMDMs together with BMDMs differentiated under 5% oxygen. We expected that increased levels of succinate and lactate in CHyp-BMDMs could give us a clue to lysosomal dysfunction because succinate has been reported as a metabolite inducing the proinflammatory status of macrophages^10,18^ and because hypoxia-induced lactate accumulation mediates macrophage polarization through histone modification^19^. However, neither succinate nor lactate increased in CHyp-BMDMs, and rather, the abundance of succinate after LPS treatment incrementally decreased according to the oxygen tension (Extended Data Fig. 5a).

As comprehensive evaluation of how prolonged hypoxia influences the LPS-induced changes in cellular metabolites, a volcano plot was drawn to determine differential metabolites in CHyp-BMDMs vs. Norm-BMDMs during LPS treatment (Fig. 4a). Out of 5 metabolites of statistical significance, pyridoxal, and pyridoxal 5’-phosphate (PLP) were dramatically decreased in CHyp-BMDMs (Fig. 4a) and incrementally decreased according to the oxygen tension (Fig. 4b). In good contrast, pyridoxine, which is contained in the culture medium as the main source of pyridoxal and PLP (Extended Data Fig. 5b), showed no changes among all oxygen conditions (Fig. 4b). The PLP reduction was also observed in the serum of a prolonged hypoxia mouse model ISAM (Fig. 4c) and in the lung tissues of mice exposed to hypoxia for 3 days (Fig. 4d). Thus, reduction in the active vitamin B6 occurred not only in macrophages but also *in vivo* under prolonged hypoxia.

**Fig. 4:**
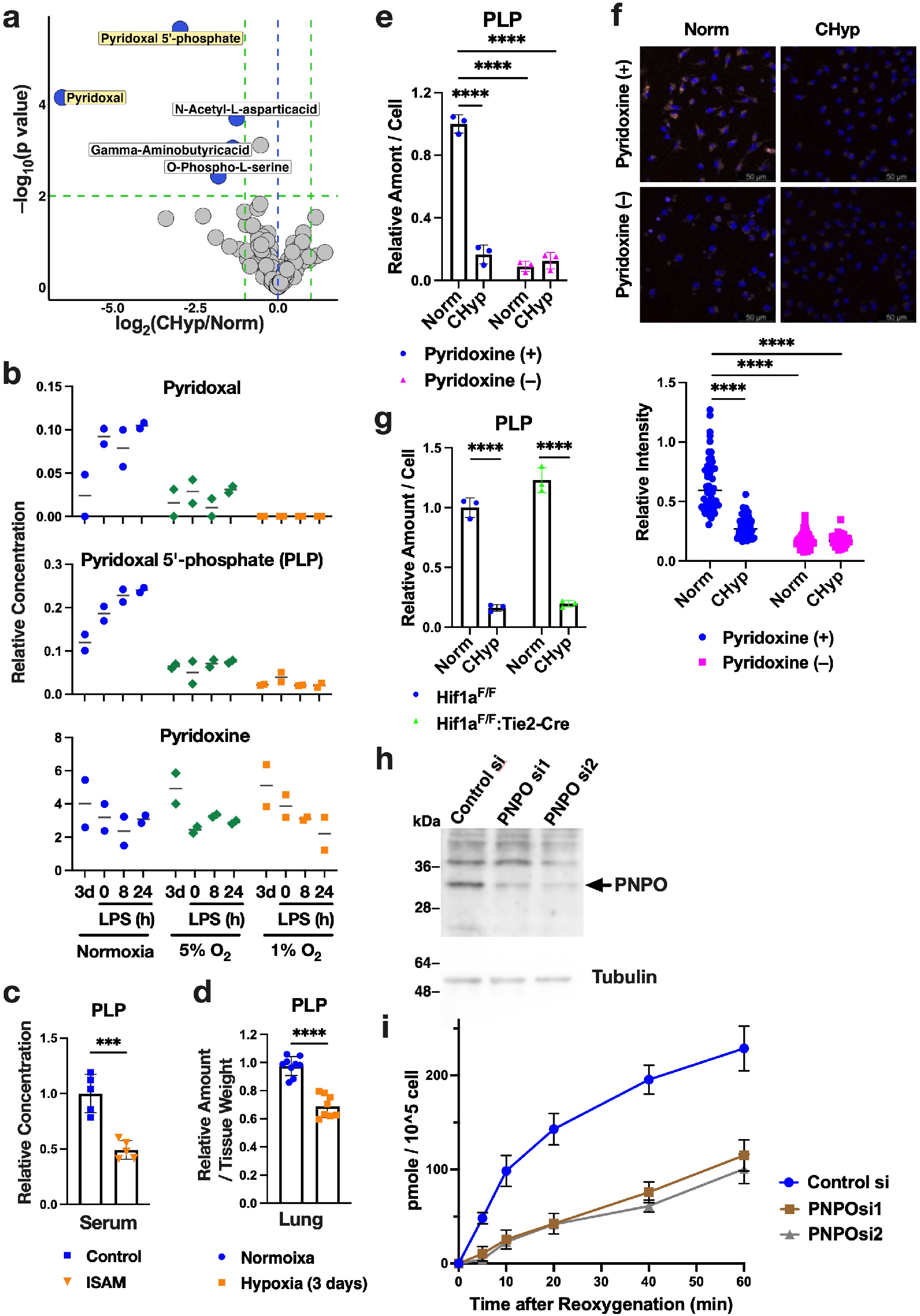
Prolonged hypoxia reduces active vitamin B6. **a**. Volcano plot showing metabolome comparison between CHyp-BMDMs and Norm-BMDMs. A horizontal dashed line indicates a significance threshold (p = 0.01), and green vertical dashed lines indicate levels of 2-fold increase and decrease. Welch’s *t* test was conducted for calculating P values. **b**. Relative amounts of pyridoxal, pyridoxal 5’-phosphate, and pyridoxine in BMDMs differentiated and stimulated with LPS under normoxia, 5% and 1% oxygen (n = 2). The metabolites were also measured on day 3 of differentiation. **c**. Relative serum concentration of pyridoxal 5’-phosphate (PLP) in ISAM and control mice (n = 5). The average value of control mice was set as 1. **d**. Relative amount of PLP in lung tissues of mice with or without hypoxia exposure for 3 days (n = 9 for 20% O_2_, n = 8 for 7% O_2_). The average value of mice in normoxia was set as 1. **e**. Relative PLP amount in CHyp-BMDMs and Norm-BMDMs with or without pyridoxine in the culture medium. The average value of Norm-BMDMs with pyridoxine was set as 1. **f**. Lysosomal acidification in CHyp-BMDMs and Norm-BMDMs with or without pyridoxine in the culture medium. Scale bars correspond to 50 μm. Representative AcidiFluor ORANGE staining (top) and its quantification (bottom). **g**. Relative PLP amount in CHyp-BMDMs and Norm-BMDMs from *Hif1a*^F/F^ mice and *Hif1a*^F/F^:Tie2-Cre mice. The average value of Norm-BMDMs from *Hif1a*^F/F^ mice was set as 1. **h**. Immunoblot analysis detecting PNPO in U937 cells treated with PNPO siRNAs or control siRNA. Tubulin was detected as a loading control. **i**. PLP increase in 1% oxygen-pre-exposed U937 cells. PLP increase upon reoxygenation is plotted. Error bars represent S.E.M. (c-e, g, i). Student’s *t*-test (c, d) and two-way ANOVA was conducted to evaluate statistical significance (e-g). ***P < 0.001, ****P < 0.0001.

To examine whether PLP is required for the maintenance of the lysosomal activity, we cultured BMDMs in the medium without pyridoxine and examined the lysosomal acidification. Pyridoxine restriction reduced the cellular PLP and inhibited the lysosomal acidification in Norm-BMDMs to the extent comparable to CHyp-BMDMs (Fig. 4e and 4f). Similar results were obtained for PMA-treated U937 cells (Extended Data Fig. 5c). These results suggest that PLP bioavailability is a key to the maintenance of the lysosomal activity.

We next addressed a molecular mechanism regulating the PLP bioavailability in response to prolonged hypoxia. Recent report described that HIF1 upregulates PDXP (see Extended Data Fig. 5b), resulting in the enhanced degradation of PLP^20^. To evaluate the HIF pathway involvement in the PLP decrease in prolonged hypoxia, we measured cellular PLP in *Hif1a*-deficient BMDMs. Prolonged hypoxia decreased PLP irrespective of the *Hif1a* status (Fig. 4g), indicating that PLP decrease induced by prolonged hypoxia is independent of HIF activity. We then focused on the synthesis of PLP from pyridoxine, which is catalyzed by PDXK and PNPO (Extended Data Fig. 5b). Because PNPO requires oxygen as a substrate, continuous hypoxic condition was expected to block the reaction catalyzed by PNPO resulting in the depletion of PLP and pyridoxal. We examined PNPO contribution to the oxygen-dependent PLP synthesis using 1% oxygen pre-exposed U937 cells with or without PNPO knockdown (Fig. 4h). The PLP increase upon reoxygenation was blunted in PNPO-knockdown cells (Fig. 4i), indicating that PNPO serves as an oxygen sensor regulating PLP bioavailability.

### Prolonged hypoxia-induced lysosomal inhibition reduces cellular Fe^2+^ and abrogates TET2 protein accumulation in response to LPS

We next investigated a molecular mechanism linking the lysosomal inhibition and proinflammatory phenotypes of macrophages under prolonged hypoxia. Because a recent study demonstrated that lysosomes are organelles that regulate cellular ferrous iron (Fe^2+^) levels and that their dysfunction causes cellular Fe^2+^ deficiency^21^, we expected that prolonged hypoxia reduced Fe^2+^ availability due to the lysosomal inhibition. Indeed, an intracellular level of Fe^2+^ detected by a fluorescent probe was remarkably decreased in CHyp-BMDMs irrespective of LPS treatment (Fig. 5a).

**Fig. 5:**
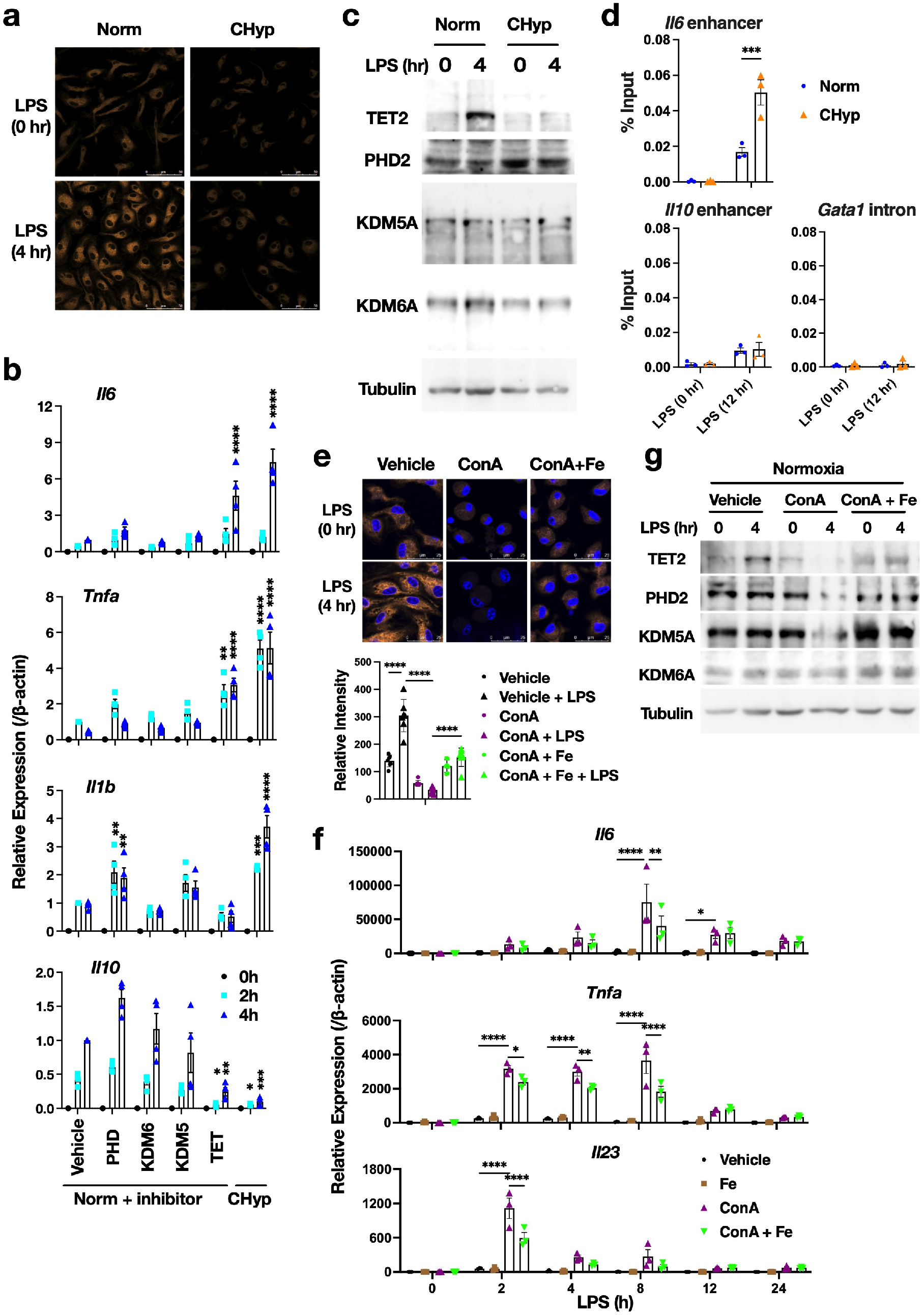
Prolonged hypoxia reduces Fe^2+^ availability and abrogates LPS-induced TET2 protein accumulation. **a**. Representative FerroOrange staining for intracellular Fe^2+^ in BMDMs treated with or without LPS for 4 h. Norm: BMDMs differentiated and stimulated with LPS under normoxia, CHyp: BMDMs differentiated and stimulated with LPS under 1% oxygen. Scale bars correspond to 50 μm. **b**. Expression of cytokine genes in BMDMs after LPS stimulation (n = 4). Norm + inhibitor: BMDMs differentiated under normoxia in the presence of 2OG-dependent dioxygenase inhibitors or vehicle (DMSO) and stimulated with LPS under normoxia. **c**. Immunoblot analysis detecting 2OG-dependent dioxygenases in BMDMs treated with or without LPS for 4 h. Tubulin was detected as a loading control. **d**. H3K27ac deposition at the *Il6* enhancer and *Il10* enhancer in BMDMs treated with or without LPS for 12 h (n = 3). The *GATA1* intron region was evaluated as a negative control locus. **e**. FerroOrange staining for intracellular Fe^2+^ in BMDMs treated with or without LPS for 4 h. BMDMs differentiated under normoxia were treated with 10 nM concanamycin A (ConA) or vehicle (DMSO) with or without 0.1 mg/ml ferric ammonium citrate (Fe) at 16 h before LPS stimulation. Hoechst was used for nuclear staining. Representative image (top) and quantification of FerroOrange fluorescence intensities by ImageJ (bottom). Scale bars correspond to 25 μm (top). Error bars represent S.D. of 7 fields per sample (bottom). **f**. Expression of cytokine genes in BMDMs after LPS stimulation (n = 3). BMDMs differentiated under normoxia were treated with 10 nM ConA or vehicle (DMSO) with or without 0.1 mg/ml ferric ammonium citrate at 16 h before LPS stimulation. **g**. Immunoblot analysis detecting 2OG-dependent dioxygenases in BMDMs treated with or without LPS for 4 h. BMDMs differentiated under normoxia were treated with 10 nM ConA or vehicle (DMSO) with or without 0.1 mg/ml ferric ammonium citrate at 16 h before LPS stimulation. Error bars represent S.E.M. of 3 or 4 biological replicates (b, d, f). Two-way ANOVA was conducted to evaluate statistical significance (b, d-f). *P < 0.05, **P < 0.01, ***P < 0.001, ****P < 0.0001. Comparisons were made against Norm + vehicle (b).

Among proteins that require Fe^2+^ for their activities, we focused on 2OG-dependent dioxygenases because they were reported to serve as oxygen sensors by switching off their activities under hypoxic conditions^3,8,9^, expecting that not only oxygen but also Fe^2+^ availability could regulated their enzymatic activities and that decline of their enzymatic activities due to Fe^2+^ limitation was responsible for the proinflammatory phenotypes of CHyp-BMDMs. To test this possibility, BMDMs were differentiated under nomoxia in the presence of inhibitors against 2OG-dependent dioxygenases, PHD, KDM6A, KDM5A and TET, and stimulated with LPS (Fig. 5b). We found that the TET inhibitor mimicked the effects of prolonged hypoxia on the gene expression of *Il6, Tnfa*, and *Il10*. Hyperactivation of *Il1b* in CHyp-BMDMs was mimicked by PHD inhibition but not by TET inhibition, which is consistent with a previous report that HIF1α induces *Il1b* expression^12^. As TET2 turned out to be an isoform that was mainly expressed in BMDMs and transcriptionally induced by LPS treatment in both Norm-BMDMs and CHyp-BMDMs (Extended Data Fig. 6a), we focused on TET2 for analysis of TET protein function hereafter.

**Fig. 6:**
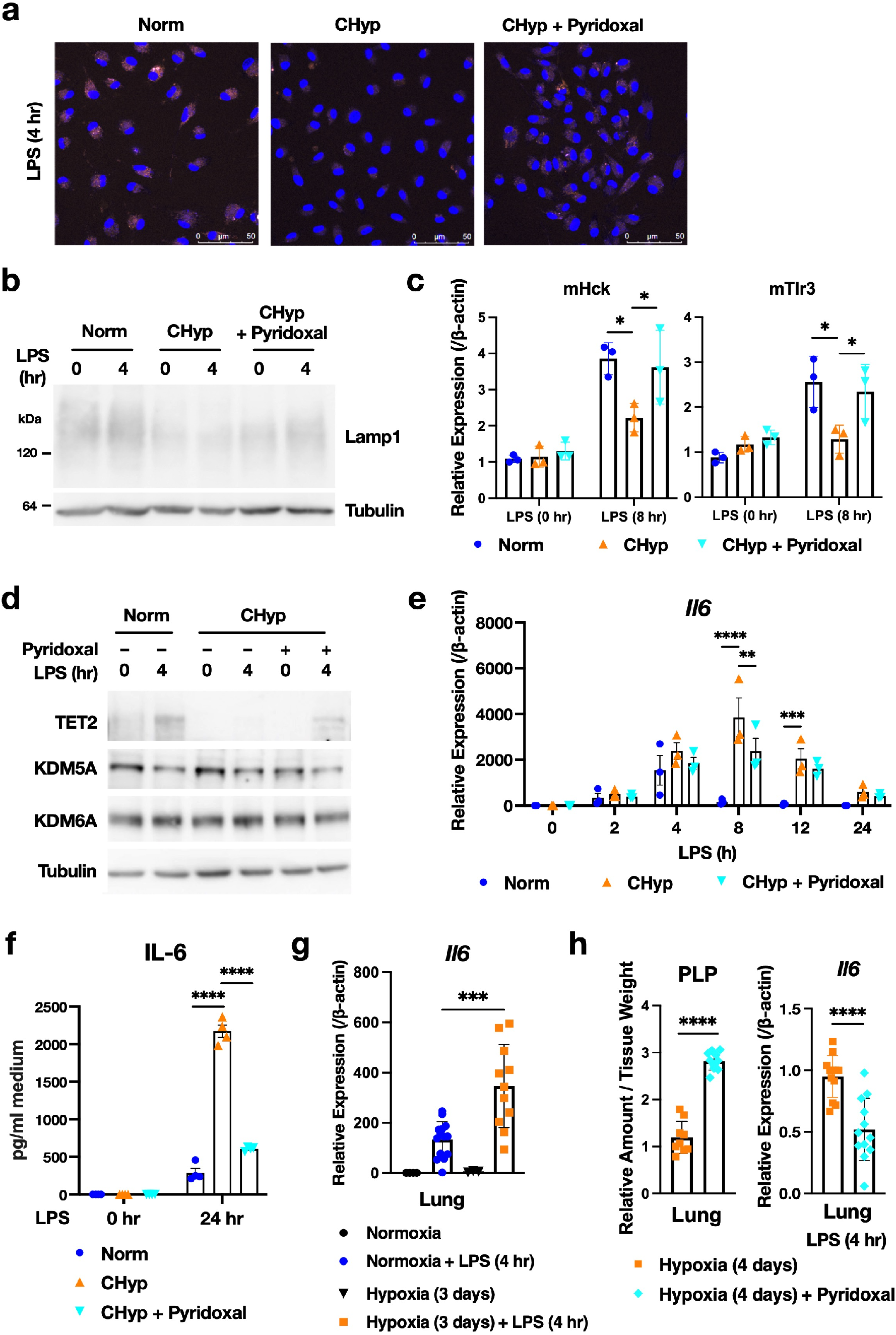
Pyridoxal supplementation restores lysosomal activity of macrophages and reverses enhanced inflammatory response in prolonged hypoxia. **a**. Representative AcidiFluor ORANGE staining for lysosomal acidification in BMDMs at 4 h after LPS treatment. CHyp + pyridoxal: BMDMs differentiated under 1% oxygen in the presence of 50 μg/ml pyridoxal. Scale bars correspond to 50 μm. **b**. Immunoblot analysis detecting Lamp1 in BMDMs treated with or without LPS for 4 h. Tubulin was detected as a loading control. **c**. Expression of representative lysosome-related genes at 8 h after LPS stimulation. **d**. Immunoblot analysis detecting 2OG-dependent dioxygenases in BMDMs treated with or without LPS for 4 h. Tubulin was detected as a loading control. **e**. Relative *Il6* expression in BMDMs after LPS stimulation (n = 3). **f**. ELISA of IL-6 in the culture supernatant of BMDMs treated with or without LPS for 24 h (n = 4). **g**. *Il6* expression at 4 h after LPS treatment in lung tissues of mice with or without hypoxia exposure for 3 days (n = 4 for normoxia, n = 14 for normoxia with LPS, n = 3 for 7% oxygen, n = 11 for 7% oxygen with LPS). **h**. Pyridoxal supplementation to mice with hypoxia exposure. Left: relative amount of PLP in lung tissues of mice exposed to hypoxia for 4 days with or without pyridoxal supplementation (n = 9 for control, n = 10 for pyridoxal supplementation). Right: *Il6* expression at 4 h after LPS treatment in lung tissues of mice exposed to hypoxia for 4 days with or without pyridoxal supplementation (n = 12). Error bars represent S.E.M. of 3-12 biological replicates (c, e, f-h). Two-way ANOVA (c, e-g) and Student’s *t*-test (h) were conducted to evaluate statistical significance. *P < 0.05, **P < 0.01, ***P < 0.001, ****P < 0.0001.

We checked the protein amount of TET2 together with PHD2, KDM5A, and KDM6A in BMDMs with or without LPS treatment (Fig. 5c). TET2 protein robustly accumulated in response to LPS in Norm-BMDMs in accordance with the increase in its mRNA, but to our surprise, the protein accumulation was almost abrogated in CHyp-BMDMs despite the LPS-induced mRNA increase (Fig. 5c and Extended Data 6a). Because TET proteins possess a high K_m_ value for Fe^2+^ compared with other 2OG-dependent dioxygenases^10^, TET proteins may be destabilized by losing Fe^2+^ under the prolonged hypoxia where Fe^2+^ availability is limited.

TET2 inactivation has been considered to underlie the proinflammatory milieu, as *Tet2* knockout mice are susceptible to DSS-induced colitis with hyperactivation of *Il6* during inflammation^22^. Recognizing that TET2 mediates DNA demethylation, we compared the LPS-induced methylome between Norm-BMDMs and CHyp-BMDMs but could not find any significant differences (data not shown). Another role of TET2 has been reported to recruit HDAC for limiting deposition of acetylated histones at the *Il6* locus in response to LPS^22^. Consistently, CHyp-BMDMs mimicked *Tet2*-deficient macrophages, showing increased deposition of acetylated histone H3K27 in the *Il6* enhancer region after LPS treatment (Fig. 5d).

Consistent with a previous report^21^, lysosomal inhibition by ConA treatment reduced cellular Fe^2+^ levels in BMDMs under normoxia (Fig. 5e). The ConA treatment also remarkably increased and decreased proinflammatory and anti-inflammatory gene expression in response to LPS, respectively (Fig. 5f and Extended Data Fig. 6b), accompanied by loss of TET2 protein accumulation (Fig. 5g). Exogenous iron supplementation partially recovered cellular Fe^2+^ in BMDMs treated with ConA (Fig. 5e). Consistent with the partial recovery of Fe^2+^, the gene expression and TET2 protein levels affected by the ConA treatment were reversed but partially (Fig. 5f, 5g and Extended Data Fig. 6b). These results suggest that lysosomal inhibition caused by prolonged hypoxia reduces Fe^2+^ availability, resulting in the abrogation of LPS-induced TET2 protein accumulation and enhancement of proinflammatory gene expression.

### Pyridoxal supplementation recovers lysosomal activity and reverses enhanced inflammatory response in prolonged hypoxia

Because PLP is a biologically active form of vitamin B6 and functions as a cofactor for various enzymes, including transaminases and decarboxylases, we suspected that depletion of pyridoxal and PLP was one of the causes of lysosomal inhibition in CHyp-BMDMs. To our delight, pyridoxal supplementation reversed the attenuated lysosomal acidification and the decreased Lamp1 protein under prolonged hypoxia (Fig. 6a and 6b). LPS-induced elevation of lysosome-related genes such as mHck and mTlr3, which was hampered in CHyp-BMDMs, was also restored by pyridoxal (Fig. 6c). These data indicated that pyridoxal supplementation recovered lysosomal function in CHyp-BMDMs. Consistent with these results, pyridoxal supplementation restored LPS-induced TET2 accumulation (Fig. 6d) and canceled the overproduction of IL-6 (Fig. 6e and 6f) in CHyp-BMDMs. These results indicate that prolonged hypoxia causes proinflammatory phenotypes of macrophages by attenuating the bioactivation of vitamin B6 that is required for lysosomal function.

We then examined whether pyridoxal supplementation cancels the *in vivo* effect of prolonged hypoxia. When LPS was injected, *Il6* expression in lung was elevated in mice exposed to hypoxia compared with those bred under normoxia (Fig. 6g). Pyridoxal was administered to the mice in the hypoxic chamber using an implantable osmotic pump, which successfully increased PLP levels in their lung tissues (Fig. 6h, left panel). When LPS was injected to the mice under 7% oxygen, *Il6* expression was attenuated in mice with the pyridoxal supplementation compared with control mice (Fig. 6h, right panel).

## Discussion

In this study, we identified PNPO-PLP axis as an oxygen-sensing mechanism operative under prolonged hypoxia, which is distinct from the canonical oxygen sensing by dioxygenases. Vitamin B6 has been revealed as an oxygen-sensitive nutrient that induces metabolic rewiring in prolonged hypoxia. An emerging concept from this work is “metabolic oxygen sensing” by which prolonged hypoxia is sensed by PNPO resulting in the gradual decline of PLP-dependent metabolism and induction of a cellular state different from that induced in response to acute hypoxia, namely lysosome-inhibited state (Extended Data Fig. 7).

The K_m_ value of PNPO for O_2_ purified from rat liver has been reported to be 182 μM by using PNP as a substrate^23^. Although it is lower than those of PHD2 (250 μM)^2^ and KDM6A (200 μM)^3^, the K_m_ value of PNPO for O_2_ is almost comparable to that of KDM4A (173 μM)^24^, which is reported to be regulated by the oxygen concentration, suggesting that the oxygen concentration also regulates PNPO activity. Of note, congenital deficiency of PNPO shows early-onset neonatal encephalopathy that closely resembles hypoxic-ischemic encephalopathy^25^. Based on these reports, we consider that PNPO is an oxygen sensor and persistent PNPO inhibition under prolonged hypoxia resulted in the depletion of PLP and pyridoxal. However, how the depletion of PLP and pyridoxal inhibits lysosomal function remains unclear.

PLP insufficiency has been shown to correlate with inflammation status^26,27^, and therapeutic potential of PLP for various inflammatory diseases has been suggested^28,29^. These previous studies are consistent with our results that prolonged hypoxia inhibits bioactivation of vitamin B6 and predisposes to exacerbated inflammation. While we focused on lysosomal function in prolonged hypoxia as an action target of PLP, PLP was also reported to suppress inflammasome activation^30^. As lysosomal dysfunction has been shown to cause excessive inflammasome activation^31^, we speculate that restoration of lysosomal integrity by PLP may limit the inflammasome activation in macrophages under prolonged hypoxia in addition to the recovery of TET2-mediated transcriptional regulation for inflammation resolution.

Recent studies have shown that lysosomes are a dynamic structure that mediates the adaptation of cell metabolism to environmental cues^32^. Inflammation is one of the major consequences of lysosomal inhibition^33,34^, which may explain proinflammatory tendencies of patients suffering from systemic chronic hypoxia. Alternatively, restraining lysosomal activity preserves the quiescence and potency of hematopoietic stem cells^35^. Considering the local hypoxic environment of the bone marrow niche where the hematopoietic stem cells reside, limited production of PLP may contribute to the inhibition of lysosomal activity for the preservation of hematopoietic stem cells. Lysosomal inhibition by the PNPO-PLP axis under chronic hypoxia is likely to underlie various pathological and physiological processes.

## Mateirals and Methods

### Mice

ISAM and their control mice^14,15^ were used for experiments at 4-5 months after birth. Blood was drawn from anesthetized mice using heparinized capillary tubes (Fisher Scientific) into the microtube within 0.5 M EDTA (pH 7.4, 2 μL) and centrifuged (3,000 rpm for 15 min at 4°C) to isolate serum for metabolome analysis. For exposure to prolonged hypoxia, C57BL/6N male mice were used at 2-3 months after birth. For preparation of bone marrow derived macrophages, wild-type mice and *Hif1a*^F/F^:Tie2-Cre mice, which were all in C57BL/6 background, were used at 2-4 months after birth. *Hif1a*^F/F^ mice (B6.129-Hif1a^tm3Rsjo^/J) and Tie2-Cre mice (B6.Cg-Tg(Tek-cre)12Flv/J) were purchased from the Jackson Laboratory (Bar Harbor, Maine, USA). These mice were bred and housed under specific pathogen-free conditions with standard animal maintenance according to the regulations of *The Standards for Human Care and Use of Laboratory Animals of Tohoku University* (Tohoku University. 2007. Standards for human care and use of laboratory animals of Tohoku University. Tohoku University, Sendai, Japan.), *The Guideline at Jichi Medical University upon approval of the Use and Care of Experimental Animals Committee of Jichi Medical University* and *The Guidelines for Proper Conduct of Animal Experiments* by the Ministry of Education, Culture, Sports, Science, and Technology of Japan (Science Council of Japan, 2006; Guidelines for the proper conduct of animal experiments; Science Council of Japan, Ministry of Education, Culture, Sports, Science, and Technology of Japan, Tokyo, Japan.).

### Dextran Sulfate Sodium (DSS)-induced colitis model

Age-matched mice (5-6 months) received 3% DSS (MP Biomedicals) in drinking water *ad libitum* for six days to induce colitis. After the mice were sacrificed by cervical dislocation, the colon was subsequently dissected for analysis. Samples were fixed in Mildform 10N (Wako) at 4°C overnight and processed for paraffin block preparation or frozen in liquid nitrogen and stored at -80°C until gene expression analysis.

### Histological analysis of DSS-treated mouse colons

Paraffin sections of colons were stained with hematoxylin and eosin. Pathological alterations were analyzed and scored following criteria that were published previously^36^. Briefly, inflammatory cell infiltration and changes in intestinal architecture were rated on a scale of 0 to 3 for each. A total score was on a scale of 0 to 6 per field. Three fields per sample, each one from the proximal, intermediate, and distal colons, were independently evaluated, and the scores were summed to determine the pathological score.

### Preparation of peritoneal exudate macrophages

For the collection of peritoneal exudate macrophages, mice were intraperitoneally injected with 2 mL of 4% thioglycolate. Peritoneal cells were isolated from exudates in the peritoneal cavity 3 days after the injection, incubated for 2 h in cell culture plates, and washed with PBS. The adherent cells were used for experiments.

### Preparation of bone marrow-derived macrophages (BMDMs)

Bone marrow cells were flushed into PBS containing 3% FBS and passed through a 70 μm nylon mech cell strainer (Falcon). The obtained whole bone marrow cells were collected by centrifugation at 800 rpm for 5 min. The cell pellet was resuspended in red cell lysis buffer (150 mM NH_4_Cl, 1 mM KHCO_3_, 0.1 mM Na_2_EDTA, 10 mM phosphate buffer) and incubated on ice for 20 min to lyse erythrocytes. After another centrifugation at 800 rpm for 5 min, cells were washed in Dulbecco’s modified Eagle’s medium (DMEM) (low-glucose with L-glutamine) once and seeded in DMEM supplemented with 40 ng/ml M-CSF (PeproTech), 10% FBS and penicillin/streptomycin. The cells were cultured at 37°C under 5% CO_2_ and saturated humidity in 1% O_2_, 5% O_2_, or normoxia. After 1 week, the culture medium was replaced with fresh DMEM supplemented with 10% FBS and penicillin/streptomycin without M-CSF.

### Reagents

2OG-dependent dioxygenase inhibitors as follows were used; KDM6A inhibitor, GSKJ4 (4594, TOCRIS Bioscience), KDM5A inhibitor, KDM5-C70 (M60192-2s, Xcess Biosciences), TET inhibitor, Bobcat339 (V4331, InvivoChem), and PHD inhibitor, GSK360A (G797600, Toronto Research Chemicals). Lysosomal inhibitors as follows were used; concanamycin A (BVT-0237-M001, AdipoGen life sciences) and bafilomycin A (BVT-0252-M001, AdipoGen Life Sciences).

### LPS treatment of macrophages

BMDMs were stimulated with 100 ng/mL LPS from Escherichia coli 0111:B4 (Sigma-Aldrich) one day after the medium change. For the normoxia control, BMDMs were differentiated in normoxia and continuously cultured in normoxia after LPS stimulation (Norm-BMDMs). For the assay of chronic hypoxia, BMDMs were differentiated in 1% O_2_ and continuously cultured in 1% O_2_ after LPS stimulation (CHyp-BMDMs). For the assay of acute hypoxia, BMDMs were differentiated in normoxia, incubated in 1% O_2_ for 12 h before LPS stimulation, and cultured in 1% O_2_ after LPS stimulation (AHyp-BMDMs). BMDMs were harvested for RNA-purification, immunoblot analysis, and iron and lysosomal staining at indicated time points.

For pretreatment with lysosomal inhibitors, BMDMs differentiated under normoxia were treated with 10 nM concanamycin A (ConA), 10 nM bafilomycin A (Baf) or vehicle (DMSO) at 16 h before LPS stimulation.

For pretreatment with inhibitors of 2OG-dependent dioxygenases, BMDMs were differentiated under normoxia in the presence of the 2OG-dependent dioxygenase inhibitors or vehicle (DMSO) and stimulated with LPS under normoxia. 2 μM GSKJ4 for KDM6A inhibition, 1 μM KDM5-C70 for KDM5A inhibition, 100 μM Bobcat339 for TET inhibition, and 5 μM GSK360A for PHD inhibition were used.

For pretreatment with pyridoxal, BMDMs were differentiated under 1% O_2_ in the presence of 50 μg/ml pyridoxal.

### RNA-seq analysis

For RNA-seq analysis of BMDMs cultured under different oxygen tension, total RNA was extracted from BMDMs using the RNeasy Mini Kit (Qiagen) in biological duplicates. Total RNA from the BMDMs was used to prepare cDNA sequencing libraries using the SureSelect Strand-Specific RNA library preparation kit (Agilent Technologies) after the polyA selection step. The libraries were sequenced on a HiSeq 2500 sequencing system (Illumina), generating 76-base single-end reads. Raw fastq sequencing files were analyzed by FastQC version 0.11.5 (available at http://www.bioinformatics.babraham.ac.uk/projects/fastqc/) to check their sequence quality, and possible adapters, poly-A tails, and low sequence quality bases were trimmed by Cutadapt version 1.15^37^. After trimming, all the remaining reads were aligned to the mm9 reference genome using STAR version 2.5.3a^38^ with the primary genome annotation in GENCODE Release M1^39^. After mapping, Cuffquant and Cuffnorm softwares, part of Cufflinks suite version 2.2.1^40^, were used to calculate an FPKM (fragments per kilobase of transcript sequence per million mapped fragments) value for each gene.

For RNA-seq analysis of BMDMs treated with lysosomal inhibitors, total RNA was extracted from BMDMs using the RNeasy Mini Kit (Qiagen) in biological duplicates. Total RNA from the BMDMs was used to prepare cDNA sequencing libraries using the TruSeq Stranded mRNA Library Prep Kit (Illumina). The libraries were sequenced on a NovaSeq 6000 sequencing system (Illumina), generating 101-base paired-end reads. Raw fastq sequencing files were analyzed as described above.

### Informatic analysis of RNA-seq data

Scatter plots and heatmaps were visualized by Rstudio packages ggplot2 and pheatmap, respectively. Enrichment analysis was performed using a browser platform of Enrichr (https://maayanlab.cloud/Enrichr/) and downloaded result file were visualized as dotplot using ggplot2. Rstudio were utilized under Rstudio version 2021.09.1+372. GSEA was conducted using a browser platform GSEA version 4.1.0.

For generating scatter plots showing correlations among impacts of hypoxia and lysosomal inhibition on LPS-induced transcriptome, we first approximated total amount of transcripts over time as the area under curve (AUC) calculated from mRNA level at each time point from 0 to 24 hours after LPS addition. AUC ratios of CHyp-BMDMs vs. Norm-BMDMs and AHyp-BMDMs vs. Norm-BMDMs were plotted for each gene to compare effects of chronic and acute hypoxia. AUC ratios of ConA-treated vs. DMSO-treated BMDMs and Baf-treated BMDMs vs. DMSO-treated BMDMs were plotted for each gene to compare effects of ConA and Baf. AUC ratios of ConA-treated BMDMs vs. DMSO-treated BMDMs and CHyp-BMDMs vs. Norm-BMDMs were plotted for each gene to compare effects of ConA and chronic hypoxia.

Gene sets for GSEA were defined as follows. ConA_up and ConA_down: upregulated and downregulated genes by more than 4-fold (log_2_ 4) by ConA treatment, respectively. Baf_up and Baf_down: upregulated and downregulated genes by more than 4-fold (log_2_ 4) by Baf treatment, respectively. CHyp_up and CHyp_down: upregulated and downregulated genes by more than 4-fold (log_2_ 4) by chronic hypoxia.

### Cell culture

U937 cells were cultured and maintained in DMEM containing 10% fetal bovine serum (FBS; Biosera, Kansas, MO, USA) under 5% CO_2_ at 37°C. U937 cells were used after differentiation into macrophage-like cells by the treatment with 10 ng /ml phorbol 12-myristate 13-acetate (PMA) for 3 days.

### Transfection of siRNA

U937 cells was transfected with siRNAs by using GenomONE-Si (Ishihara Sangyo) according to the manufacturer’s protocol. PMA was added right after the transfection. The cells were analyzed 72 hours after the transfection. Control (MISSION® siRNA Universal Negative Control #1,Sigma), PNPOsi1(SASI_Hs01_00068079, Sigma), and PNPOsi2(SASI_Hs02_00351341,Sigma) were used.

To prepare the cells for PLP measurement, U937 cells treated with siRNA and PMA were incubated in 1% O_2_ for 72 hours. After being washed in PBS once, the cells were reoxygenated in normoxia-equilibrated and prewarmed medium containing 4.0 mg/L pyridoxine hydrochloride.

### ChIP assay

ChIP assays were performed with BMDMs differentiated in 1% O_2_ and normoxia using anti-H3K27ac antibody (MABI0309, MAB Institute). The cells were treated with 100 ng/ml LPS for 12 h and cross-linked with 1% formaldehyde for 10 min. The samples were then lysed and sonicated to shear DNA. Sonication was conducted according to previously described procedures^41,42^. The solubilized chromatin fraction was incubated overnight with anti-H3K27ac antibody that was prebound to Dynabeads anti-rabbit IgG (Life Technologies). Precipitated DNA was analyzed by real-time PCR. The primer sets used in the ChIP assay are listed in Extended Data Table 1.

### ELISA

Peritoneal macrophages or BMDMs at 2 × 10^6^ cells/mL were incubated with 100 ng/mL LPS for 12 h or 24 h to measure TNF-α and IL-6 and further incubated for 2 h with 1 mM ATP to measure IL-1β. The culture supernatants were assessed for the cytokines using mouse TNF-α, IL-6, and IL-1β ELISA kits (R&D Systems).

### RNA purification and quantitative RT-PCR

Total RNA samples were prepared from cells and tissues using ISOGEN (Nippon Gene) or ReliaPrep™ RNA Miniprep Systems (Promega) according to the manufacturer’s instructions. First-strand cDNA was synthesized from 100 ng of total RNA using ReverTra Ace qPCR RT Master Mix with gDNA Remover (TOYOBO). Real-time PCR was performed in triplicate for each sample with QuantStudio real-time PCR system (Thermo Fisher Scientific, Waltham, MA, USA) using KAPA SYBR FAST qPCR Master Mix (Kapa Biosystems, Wilmington, MA, USA). Expression levels of *Actb* (beta-actin) gene were used as internal controls for normalization. The primer sets used in the real-time PCR are listed in Extended Data Table 2.

### Immunoblot analysis

Immunoblot analyses were performed as described previously^43^. The antibodies that were used were anti-TET2 (ab124297, Abcam), KDM5a (ab70892, Abcam), anti-KDM6a (33510S, Cell Signaling), anti-PHD2 (NB100-2219, Novus), anti-Lamp1 (ab25245, Abcam), anti-PNPO (15552-1-AP, Proteintech), and anti-Tubulin (T9026, Sigma).

### Detection of lysosomal acidification

Lysosomal acidity was observed by using AcidiFluor ORANGE kit (Goryo Chemical) according to the manufacturer’s instructions. Nuclei were imaged with Hoechst 33342 (Dojindo). Briefly, BMDMs and U937 cells were prepared in 4-chamber 35 mm culture dishes at a density of 2×10^5^ cells/well under the indicated conditions. The cells were stained with 2 μM AcidiFluor ORANGE and 0.5 μg/mL Hoechst 33342 for 2 h at 37°C under 5% CO_2_. Then, the cells were washed three times with PBS and observed with confocal microscopy (TCS SP8, Leica). AcidiFluor ORANGE was detected at Ex/Em = 552/570-590 nm. The fluorescence intensities were quantified by using LASX software (Leica). A single cell was circled, and the intensity of each circle was quantified. The AcidiFluor ORANGE intensity was normalized by Hoechst 33342 intensity for each cell.

### Detection of intracellular ferrous iron

The iron concentration was assessed using FerroOrange (DojinDo) to measure intracellular ferrous iron levels according to the manufacturer’s protocol. Briefly, BMDMs were prepared in 4-chamber 35 mm culture dishes at a density of 2×10^5^ cells/well and treated with or without LPS for 4 h. The cells were washed three times with PBS and stained with FerroOrange working solution for 30 min at 37°C under 5% CO_2_. Then, the cells were observed with confocal microscopy (TCS SP8, Leica) at Ex/Em = 552/561-570 nm. The fluorescence intensity was measured by ImageJ software (National Institute of Health).

### Treatment with ferric ammonium citrate

BMDMs differentiated under normoxia were treated with 10 nM concanamycin A (ConA) or vehicle (DMSO) with or without 0.1 mg/ml ferric ammonium citrate (Fe) at 16 h before LPS stimulation. Then cells were harvested for ferrous iron detection, gene expression analysis and immunoblot analysis.

### Methylome analysis

Methylome data were obtained with whole-genome bisulfite sequencing (WGBS). Libraries for WGBS were prepared based on post-bisulfite adaptor tagging (PBAT)^44^ using an improved protocol tPBAT^45^. One hundred fifty nanograms of purified genomic DNA was used for each sample. PCR amplification was not performed for any library. After the preparation, libraries of three biological replicates tagged with different indices were mixed equally. The mixture was sequenced using the HiSeq X ten system. Sequencing was performed by Macrogen Inc. (Tokyo, Japan). The obtained reads were mapped to the reference genome and summarized with an in-house pipeline^44^.

### Metabolome analysis

Total metabolites were extracted from BMDMs and mouse serum according to the Bligh and Dyer method with minor modifications^46^. For metabolite measurement in BMDMs, BMDMs were washed twice with cold PBS and harvested in 1 mL cold methanol (–30°C) containing 10-camphorsulfonic acid (1.5 nmol) and piperazine-1,4-bis(2-ethanesulfonic acid) (PIPES) (1.5 nmol) as internal standards (ISs). The samples were vigorously mixed by vortexing for 1 min followed by 5 min of sonication. To precipitate protein, the methanol extracts were incubated on ice for 5 min. The extracts were then centrifuged at 16,000 g for 5 min at 4°C, and the resultant supernatant was collected. Protein concentrations in the pellet were determined using a Pierce™ BCA Protein Assay Kit (Thermo Fisher Scientific). Then, 600 μL of supernatant was transferred to another tube, and 600 μL of chloroform and 480 μL of water were added. After centrifugation at 16,000 *g* and 4°C for 5 min, 800 μL of the upper layer was isolated and used for hydrophilic metabolite analysis.

For metabolite measurement in mouse serum, 50 μL of serum was mixed with 940 μL of methanol and 10 μL of IS solution containing 10-camphorsulfonic acid (2.0 nmol) and PIPES (2.0 nmol). The samples were centrifuged at 16,000 g at 4°C for 5 min, and the supernatant (400 μL) was collected in clean tubes. After mixing with 400 μL of chloroform and 320 μL of water, phase separation of aqueous and organic layers was performed via centrifugation (16,000 *g*, 4°C, 5 min). The aqueous (upper) layer (500 μL) was transferred into a clean tube. After the aqueous layer extracts were evaporated under vacuum, the dried extracts were stored at −80°C until the analysis of hydrophilic metabolites. Prior to analysis, the dried aqueous layer was reconstituted in 50 μL of water.

Anionic polar metabolites (e.g., succinate, lactate, PLP) were analyzed via ion chromatography (Dionex ICS-5000^+^ HPIC system, Thermo Fisher Scientific) with a Dionex IonPac AG11-HC-4 μm guard column (2 mm i.d. × 50 mm, 4 μm particle size, Thermo Fisher Scientific) and a Dionex IonPac AS11-HC-4 μm column (2 mm i.d. × 250 mm, 4 μm particle size, Thermo Fisher Scientific) coupled with a Q Exactive, high-performance benchtop quadrupole Orbitrap high-resolution tandem mass spectrometer (Thermo Fisher Scientific) (IC/HRMS/MS)^47^. Cationic polar metabolites (e.g., pyridoxal, pyridoxine) were analyzed via liquid chromatography (Nexera X2 UHPLC system, Shimadzu Co., Kyoto, Japan) with a Discovery HS F5 column (2.1 mm i.d. × 150 mm, 3 μm particle size, Merck) coupled with a Q Exactive instrument (PFPP-LC/HRMS/MS)^47^.

### Informatic analysis of metabolome data

Metabolome data obtained from BMDMs differentiated in 1% O_2_ and normoxia irrespective of the LPS treatment were used for the creation of a volcano plot. Welch’s *t* test was conducted for statistical significance.

### Quantification of PLP

PLP was quantified using VB6 Enzymatic assay kit (Bühlmann) according to the manufacture’s protocol. Briefly, BMDMs and U937 cells cultured in 24 well plates were washed with PBS once and then collected in 100ul methanol. The methanol extracts were diluted by “substrate buffer” for the measurement. Lung tissues were weighed and homogenized in methanol at the ratio of 100 mg tissue per mL methanol. After centrifugation at 5,000 x g for 5 min, the supernatant was diluted by 500-fold and added to the “substrate buffer” for the measurement.

### Pyridoxine restriction culture of BMDM and U937 cells

Pyridoxine-depleted DMEM was manufactured on request (Cell Science & Technology Institute Inc.). For experiments using BMDM, the pyridoxine-depleted DMEM was used from the beginning of the BMDM differentiation. For experiments using U937, normal DMEM was changed to the pyridoxine-depleted DMEM at the time of PMA addition.

### Hypoxia exposure and LPS treatment of mice

A hypoxia chamber, in which the oxygen concentration was regulated by an oxygen controller (ProOx; BioSpherix) with a nitrogen generator (Nilox; Sanyo Electronic Industries), was used to expose the mice to hypoxic conditions. To avoid accumulation of moisture and carbon dioxide, a moisture absorber and Litholyme® were set in the chamber, respectively. To reduce ammonia during the hypoxia exposure, cat litter was layered beneath the wooden chip bedding of mouse cages. Mice were exposed to 7% O_2_ for 3 days and sacrificed for PLP measurement. For LPS treatment, mice were exposed to 7% O_2_ for 3 days and given intraperitoneal injection of LPS at the dose of 5 mg/kg body weight. The mice were sacrificed at 4 h after the LPS injection. For supplementation of pyridoxal, ALZET® osmotic pump 1007D containing 150 mg/mL pyridoxal hydrochloride (P9130, Sigma-Aldrich) in PBS was subcutaneously implanted to the mice under general anesthesia, and the exposure to 7% O_2_ was started from the next day. Control mice underwent sham operation of the back skin under general anesthesia and were similarly exposed to the hypoxia. After 4-day exposure to 7% O_2_, the mice were injected with LPS (5 mg/kg body weight) and sacrificed at 4 h after the treatment.

### Statistical analysis

Statical analysis was conducted using Prism 7 (GraphPad Software, CA, USA). Data are presented as means ± S.E.M. or S.D. and were tested with unpaired Student’s *t* test, Welch’s *t* test and one–way or two-way analysis of variance (ANOVA) followed by Tukey’s multiple comparison test. P value of < 0.05 was considered significant.

### Data availability

RNA-seq data were deposited at GEO (GSE181641, GSE181642, GSE181643).

## Acknowledgments

We thank the Department of Gene Expression Regulation and the Biomedical Research Core of the Institute of Development, Aging and Cancer for discussions and technical support. This work was supported by JSPS (grant numbers 17K08618 (H.S.), 20K07320 (H.S.), 17H06299 (T.B.), 17H06304 (T.B.), 20H04832 (H.M.), 21H04799 (H.M.), 21H05258 (HM) and 21H05264 (HM)), AMED under grant number JP21gm5010002 (H.M.), Japan-Sweden Research Cooperative Program between JSPS and STINT (grant number JPJSBP120195402 (H.M)), and Canon Medical Systems (K.K., H.M.). This research was also supported by the Platform Project for Supporting Drug Discovery and Life Science Research (Basis for Supporting Innovative Drug Discovery and Life Science Research (BINDS)) from AMED under Grant Number JP21am0101103 (support number 2320). The funders had no role in the study design, data collection and analysis, decision to publish or manuscript preparation.

## Author contributions

H.S. designed the study, conducted the experiments, analyzed the data and wrote early draft of the paper. H.T. conducted bioinformatic analyses and wrote the paper. A.K. conducted the experiments and analyzed the data. H.A. analyzed RNA-seq data and wrote the paper. N.O., N.K., S.K., Z.L. and K. Kato conducted the experiments. F.K. conducted the experiments and wrote the paper. F.M. and T. Ito conducted methylome analysis and wrote the paper. M.T., Y.I., and T.B. conducted metabolome analysis and wrote the paper. T. Isagawa, N.T., N.S. and M.Y. provided critical materials and analyzed and interpreted the data. H.F., and H.Y. analyzed and interpreted the data. K. Kinoshita analyzed and interpreted the data and wrote the paper. H.M. designed the study, conducted the experiments, supervised the research, analyzed the data and wrote early draft of the paper.

