## Supplementary materials for "PNPO-PLP Axis Senses Prolonged Hypoxia by Regulating Lysosomal Activity"

**This file includes:**

Extended Data Figures 1 to 7
Extended Data Tables 1 and 2

### Extended Data Figures

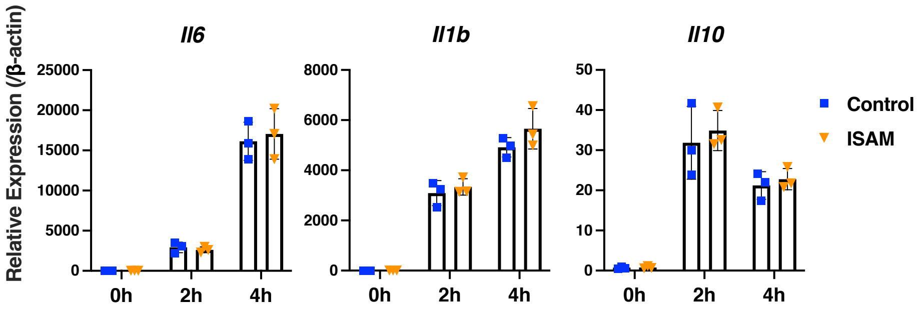


**Extended Data Fig. 1: Expression of cytokine genes in BMDMs after LPS treatment.** BMDMs of ISAM and control mice were differentiated under normoxia. Error bars represent S.E.M. of 3 biological replicates. Two-way ANOVA was conducted to evaluate statistical significance.

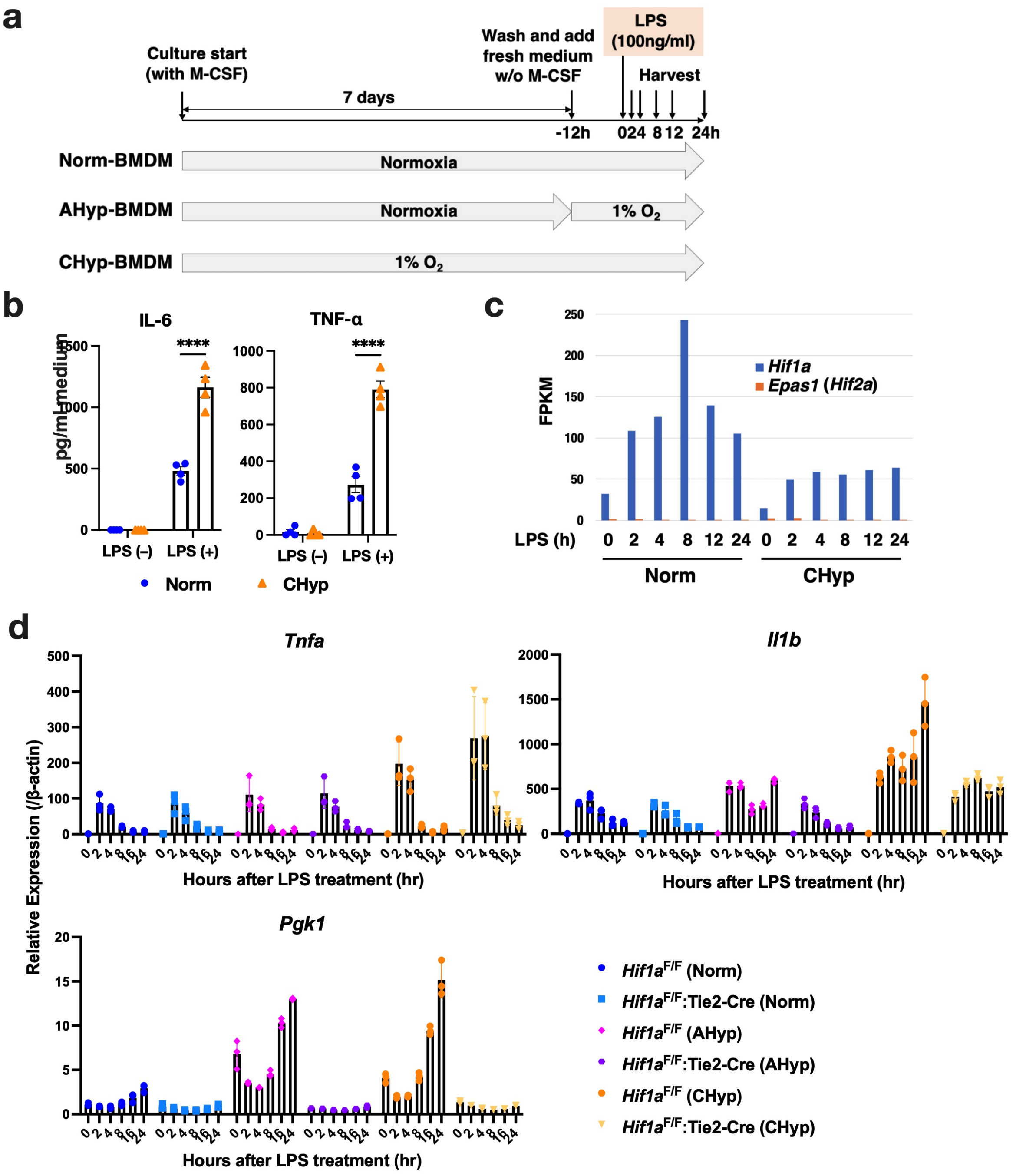


**Extended Data Fig. 2: Inflammatory response of BMDMs differentiated under different oxygen tension.**

(**a**) Experimental design for BMDMs differentiation and stimulation with LPS (100 ng/ml). BMDMs were harvested for RNA-seq analysis at 0, 2, 4, 8, 12 and 24 h after LPS treatment.

(**b**) ELISA of cytokines in culture supernatant of BMDMs stimulated with or without LPS for 12 h (n = 4). Error bars represent S.E.M. Two-way ANOVA was conducted to evaluate statistical significance. ****P < 0.0001.

(**c**) FPKM values of *Hif1a* and *Epas1*/*Hif2a* expression obtained from the RNA-seq analysis.

(**d**) Gene expression in BMDMs after LPS stimulation (n = 3). Norm-, AHyp-, and CHyp-BMDMs generated from *Hif1a*$^{F/F}$ mice and *Hif1a*$^{F/F}$:Tie2-Cre mice were examined. *Pgk1* is a target gene of HIF-1α. Error bars represent S.E.M. of 3 biological replicates.

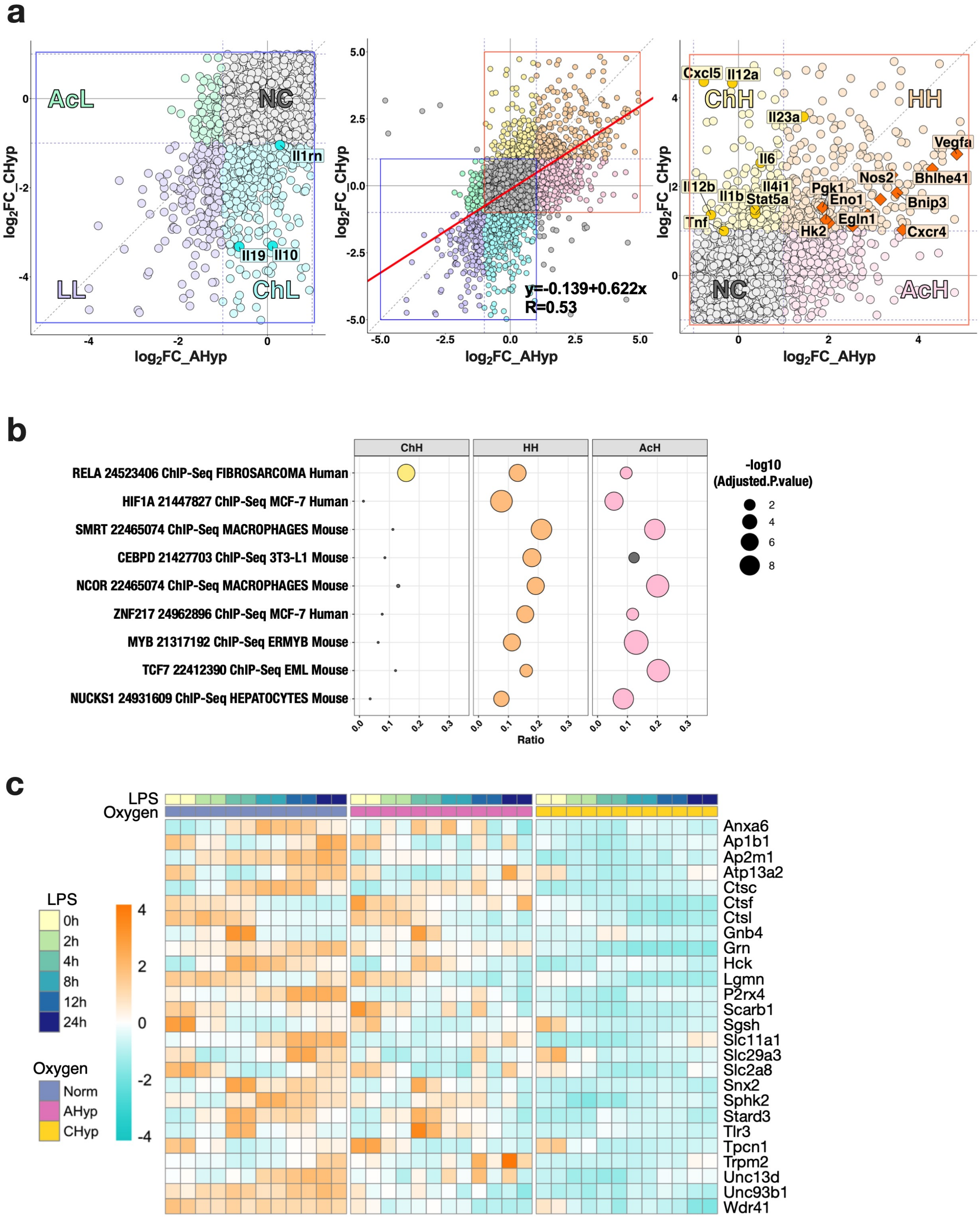


**Extended Data Fig. 3: Comparison of LPS-induced transcriptome in BMDMs differentiated under different oxygen tension.**

(**a**) Scatter plots showing a correlation of gene expression fold changes (AUC ratios) by chronic and acute hypoxia vs. normoxia in BMDMs. A horizontal axis indicates $\log_2$ fold change of

CHyp vs. Norm, and a vertical axis indicates $log_2$ fold change of AHyp vs. Norm. Areas enclosed by red and blue squares (middle panel) are highlighted as those containing upregulated and downregulated genes, respectively (right and left panels). ChH: specifically upregulated in chronic hypoxia, AcH: specifically upregulated in acute hypoxia, HH: upregulated in both conditions, ChL: specifically downregulated in chronic hypoxia, AcL: specifically downregulated in acute hypoxia, LL: commonly downregulated in both conditions, NC: not changed. The strength of the correlation was evaluated with Pearson product-moment correlation coefficient.
(**b**) Enrichr analysis (ChEA 2016) of upregulated genes. ChH, HH and AcH are gene groups indicated in panel A (right). Circle sizes indicate adjusted P values, and circle colors indicate statistical significance (yellow, orange, and pink: adjusted P value $< 0.05$, gray: not significant).
(**c**) A heatmap illustrating RNA-seq data of genes that belong to ChL class and assigned to "lysosome" in GO database. Norm: BMDMs differentiated and stimulated with LPS under normoxia, AHyp: BMDMs differentiated under normoxia and stimulated with LPS under 1% $O_2$, CHyp: BMDMs differentiated and stimulated with LPS under 1% $O_2$.

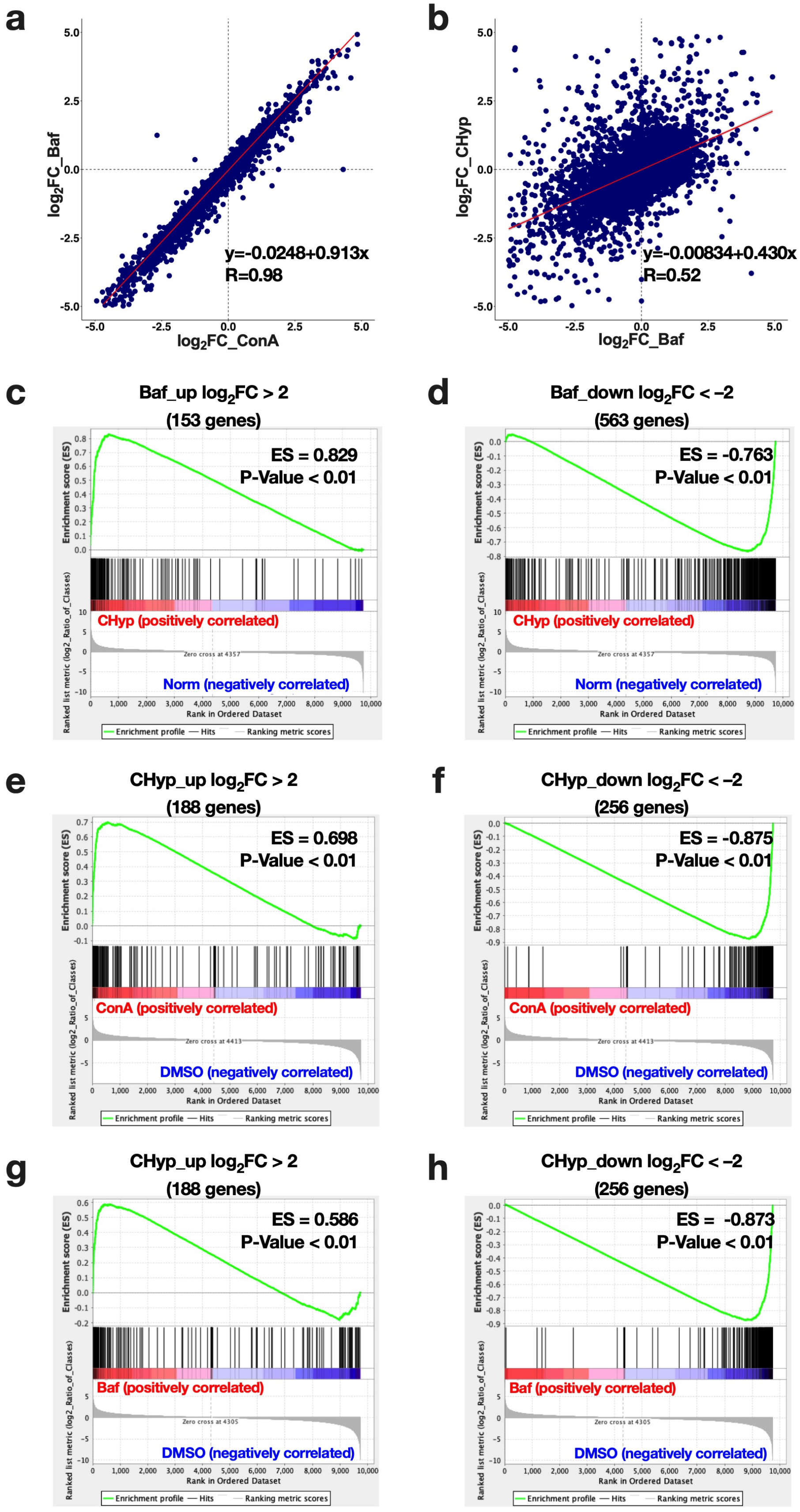
a
log2FC_Baf
log2FC_ConA
y=-0.0248+0.913x
R=0.98
b
log2FC_CHyp
log2FC_Baf
y=-0.00834+0.430x
R=0.52
c
Baf_up log2FC > 2
(153 genes)
ES = 0.829
P-Value < 0.01
CHyp (positively correlated)
Norm (negatively correlated)
d
Baf_down log2FC < –2
(563 genes)
ES = -0.763
P-Value < 0.01
CHyp (positively correlated)
Norm (negatively correlated)
e
CHyp_up log2FC > 2
(188 genes)
ES = 0.698
P-Value < 0.01
ConA (positively correlated)
DMSO (negatively correlated)
f
CHyp_down log2FC < –2
(256 genes)
ES = -0.875
P-Value < 0.01
ConA (positively correlated)
DMSO (negatively correlated)
g
CHyp_up log2FC > 2
(188 genes)
ES = 0.586
P-Value < 0.01
Baf (positively correlated)
DMSO (negatively correlated)
h
CHyp_down log2FC < –2
(256 genes)
ES = -0.873
P-Value < 0.01
Baf (positively correlated)
DMSO (negatively correlated)
Enrichment score (ES)
Ranked list metric (log2_Ratio_of_Classes)
Rank in Ordered Dataset
Enrichment profile
Hits
Ranking metric scores

**Extended Data Fig. 4: Comparison between impacts of prolonged hypoxia and lysosomal inhibition on the LPS-induced transcriptome.**
(**a, b**) Scatter plots showing correlations of gene expression fold changes (AUC ratios) by ConA and Baf (a) and by prolonged hypoxia and Baf (b) in BMDMs. For panel A, a horizontal axis indicates $\log_2$ fold change of ConA treatment vs. DMSO, and a vertical axis indicates $\log_2$ fold change of Baf treatment vs. DMSO. For panel B, a horizontal axis indicates $\log_2$ fold change of Baf treatment vs. DMSO, and a vertical axis indicates $\log_2$ fold change of CHyp vs. Norm (shown in Fig. 2c). The strength of each correlation was evaluated with Pearson product-moment correlation coefficient.
(**c, d**) Gene set enrichment analysis comparing the impacts of prolonged hypoxia with lysosomal inhibition (Baf). Gene sets were defined as upregulated (c) or downregulated (d) genes by more than 4-fold ($\log_2$ 4) by Baf treatment. Changes in the LPS-induced transcriptome by prolonged hypoxia were analyzed against the gene sets.
(**e-h**) Gene set enrichment analysis comparing the impacts of lysosomal inhibition with prolonged hypoxia. Gene sets were defined as upregulated (e, g) or downregulated (f, h) genes by more than 4-fold ($\log_2$ 4) by prolonged hypoxia. Changes in the LPS-induced transcriptome by lysosomal inhibitors, ConA (e, f) and Baf (g, h), were analyzed against the gene sets.

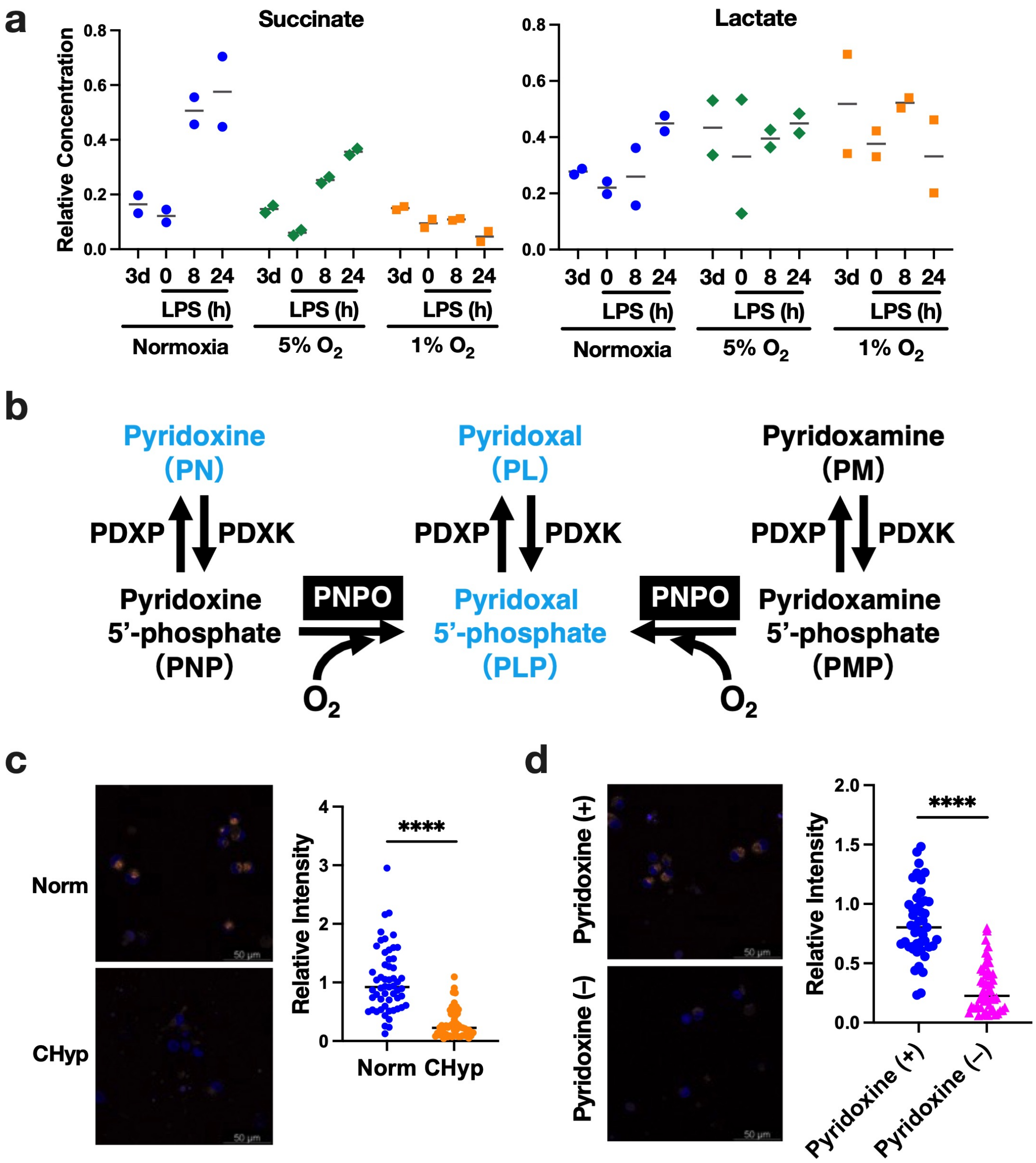


**Extended Data Fig. 5: Metabolite changes induced by prolonged hypoxia and its effect on lysosomal activity.**

(**a**) Relative amounts of cellular succinate and lactate in the BMDMs differentiated and stimulated with LPS under normoxia, 5% $O_2$ and 1% $O_2$ (n=2). The metabolites were also measured on day 3 of differentiation.

(**b**) Schematic illustration indicating vitamin B6 metabolism and catalyzing enzymes. PDXP: pyridoxal phosphatase, PDXK: pyridoxal kinase, PNPO: pyridoxine 5'-phosphate oxidase.

(**c, d**) Lysosomal acidification in PMA-treated U937 cells under different oxygen tension (c) and with or without pyridoxine in the culture medium (d). Scale bars correspond to 50 µm. Representative AcidiFluor ORANGE staining (left) and its quantification (right). Student's *t*-test was conducted to evaluate statistical significance. ****$P < 0.0001$.

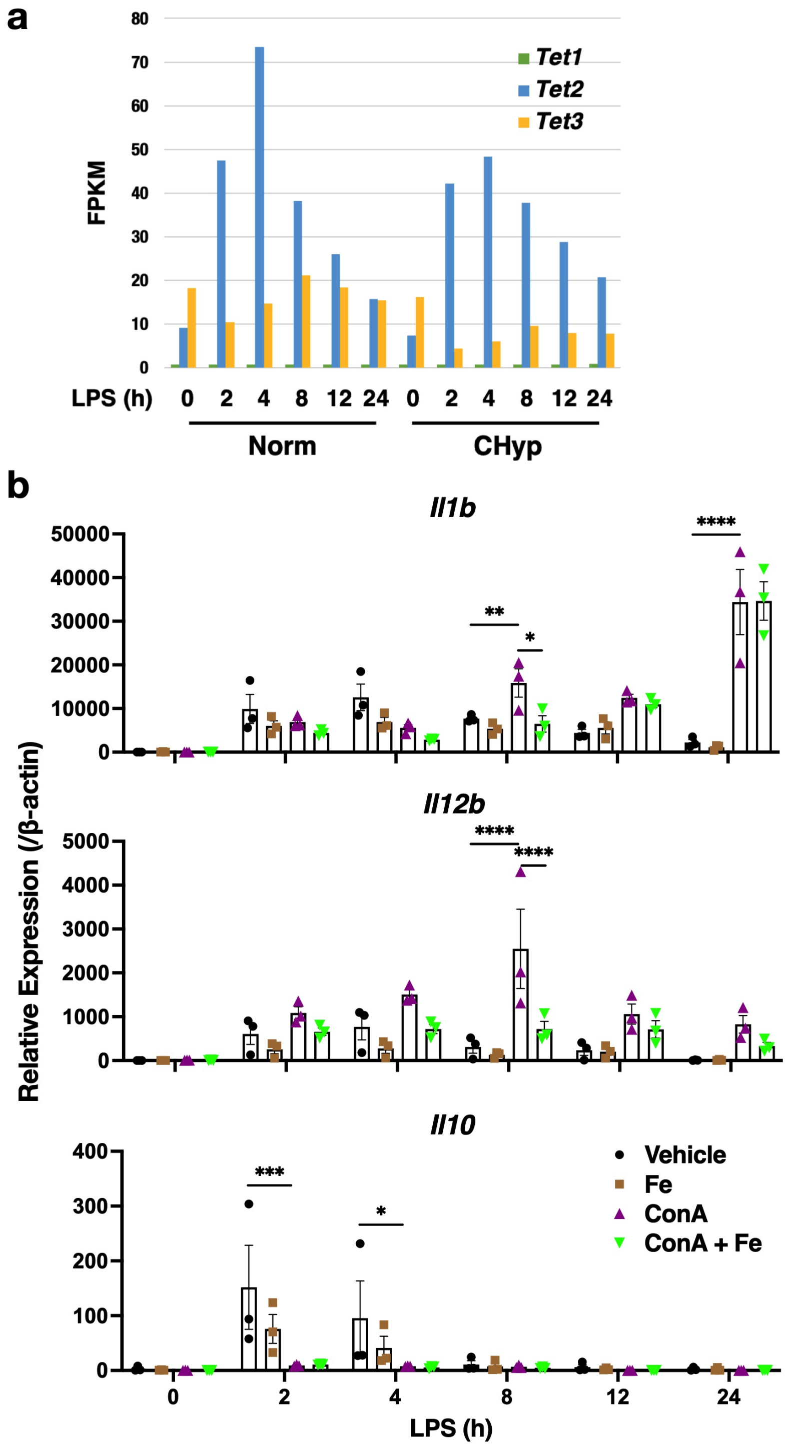


**Extended Data Fig. 6: LPS-induced gene expression in BMDM.**
(**a**) FPKM values of *Tet1*, *Tet2* and *Tet3* expression obtained from the RNA-seq analysis. Norm: BMDMs differentiated and stimulated with LPS under normoxia, CHyp: BMDMs differentiated and stimulated with LPS under 1% $O_2$.
(**b**) Expression of cytokine genes in BMDMs after LPS stimulation (n=3). BMDMs differentiated under normoxia were treated with 10 nM ConA or vehicle (DMSO) with or

without 0.1 mg/ml ferric ammonium citrate at 16 h before LPS stimulation. Error bars represent S.E.M. of 3 biological replicates. Two-way ANOVA was conducted to evaluate statistical significance. *P<0.05, **P<0.01, ***P<0.001, ****P<0.0001.

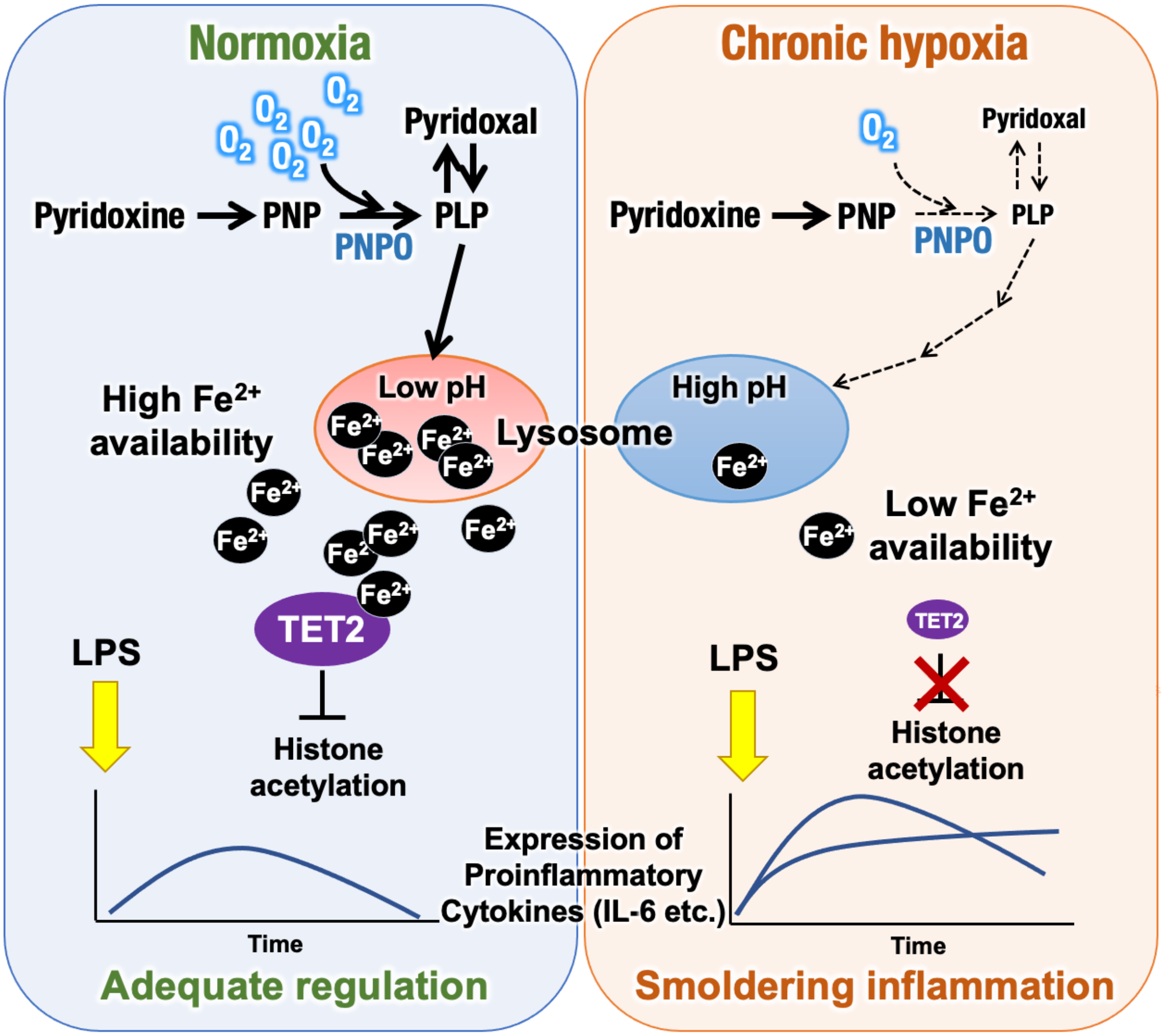


**Extended Data Fig. 7: Metabolic oxygen sensing by PNPO-PLP axis controls inflammatory response of macrophages.**
Prolonged hypoxic condition reduces PNPO activity and gradually decreases PLP that is required for the maintenance of lysosomal function. Lysosomal inhibition caused by PLP insufficiency limits $Fe^{2+}$ availability and inhibits TET2 function, resulting in the delayed resolution of inflammation.

#### Extended Data Tables

##### Extended Data Table 1: Primers Used for ChIP Assay.

| Name | Sequence (5’-3’) |
| --- | --- |
| mIl6 enhancer forward | GGGAAATCGTGGAAATGAGA |
| mIl6 enhancer reverse | CCAGCAAAGAGGTGAGAAAGA |
| mIl10 enhancer forward | GGGCTGGAGTTGTATGGAGT |
| mIl10 enhancer reverse | CACGGCAAGAGTGTGGTATG |
| mGata1 intron forward | TGTCAGGCTTCCATTTGAGA |
| mGata1 intron reverse | CCCAGATAACCTCGTGCTGT |

**Extended Data Table 2: Primers used for RT-PCR.**

| Name | Sequence (5’-3’) |
|---|---|
| mIl6 forward | CTGCAAGAGACTTCCATCCAG |
| mIl6 reverse | AGTGGTATAGACAGGTCTGTTGG |
| mIl23 forward | CTGCTTGACTCTGACATC |
| mIl23 reverse | CACTGCTGACTAGAACTC |
| mIl1β forward | TGCCACCTTTTGACAGTGATG |
| mIl1β reverse | TGATGTGCTGCTGCGAGATT |
| mIl12b forward | AGACCCTGCCCATTGAACTG |
| mIl12b reverse | GGCGGGTCTGGTTTGATGAT |
| mTnfα forward | CACGCTCTTCTGTCTACTGAA |
| mTnfα reverse | GGCTACAGGCTTGTCACTCGA |
| mIl10 forward | GGCGCTGTCATCGATTTCTC |
| mIl10 reverse | ATGGCCTTGTAGACACCTTGG |
| mBnip3 forward | GTTACCCACGAACCCCACTTT |
| mBnip3 reverse | GTGGACAGCAAGGCGAGAAT |
| mPgk1 forward | GTCGTGATGAGGGTGGACTT |
| mPgk1 reverse | GACAACGGACTTGGCTCCA |
| mHck forwward | TTGAGGTTGACGCACCTGTT |
| mHck reverse | AGCCCAGATTATGGGTGCAA |
| mTlr3 forward | GAGTACACAGCTCGGGAAGG |
| mTlr3 reverse | AGCCCAGATTATGGGTGCAA |
| mbetaActin forward | CGGTTCCGATGCCCTGAGGCTCTT |
| mbetaActin reverse | CGTCACACTTCATGATGGAATTGA |
